# Granulysin-Based pH-Sensitive Antimicrobial Nanocarriers for Treatment of Multidrug-Resistant Bacterial Wound Infections

**DOI:** 10.64898/2026.03.26.714505

**Authors:** Owais A. Hameed, Mark Gontsarik, Patricia Matthey, Oriana Coquoz, Jules Denis Pierre Valentin, Stefan Salentinig, Michael Walch

## Abstract

Multidrug resistant (MDR) bacterial wound infections are an increasing clinical challenge and require alternatives to conventional antibiotics. Although antimicrobial proteins offer promise, their therapeutic use is limited by poor stability, proteolytic degradation, reduced activity under physiological conditions, and potential toxicity. This work reports pH-sensitive lipid nanocarriers composed of granulysin (GNLY) and oleic acid (OA) for antimicrobial delivery to infected tissues. At neutral pH, GNLY is retained within OA-based nanocarriers and protected from proteolytic degradation. At pH 5.0, such as in infected wounds, the carriers undergo structural reorganization and release GNLY, restoring antimicrobial activity. OAGNLY (32 µg/mL) achieved >3-log reductions in Staphylococcus aureus and Escherichia coli within 1 hour, and up to 4-log reductions in Pseudomonas aeruginosa and Acinetobacter baumannii, at physiological salt concentrations where free GNLY was largely inactive. Minimum inhibitory concentrations were 16 µg/mL for MRSA and 32 µg/mL for colistin-resistant E. coli. Ultrastructural analysis using transmission electron microscopy revealed disruptions of bacterial membranes and intracellular structures following OAGNLY treatment. In a murine surgical wound infection model, topical application of OAGNLY for 4 hours reduced bacterial burden by >5 logs and significantly decreased inflammation, as confirmed by histological analysis. In parallel, OAGNLY demonstrated minimal cytotoxicity to mammalian cells at active concentrations. These findings identify OAGNLY nanocarriers as a promising platform for pH-responsive delivery of GNLY and highlight their potential application for treating MDR skin and soft tissue infections..

## 1 Introduction

Bacterial wound and skin/soft tissue infections represent a major clinical challenge, as they impair healing and can progress to chronic disease states.[1] Bacterial pathogens are continually evolving to acquire resistance to conventional antibiotics, complicating treatment and having tremendous implications for the global healthcare system.[2] Infections caused by multidrug-resistant (MDR) pathogens, including *methicillin-resistant Staphylococcus aureus (MRSA), carbapenem-resistant Escherichia coli*, *Pseudomonas aeruginosa,* and *Acinetobacter baumannii,* are particularly concerning in vulnerable populations, including hospitalized and immunocompromised patients, where they are associated with severe complications and high mortality.[3, 4] In recent years, colistin, the last-resort antibiotic for highly problematic Gram-negative bacteria, has faced challenges due to the emergence and rapid spread of the mcr-1 gene, causing resistance.[5–8] Together with the limited development of new antibiotics, this underscores an urgent need for alternative therapeutic strategies that can overcome current resistance mechanisms.

Antimicrobial peptides and proteins (AMPs) have attracted considerable interest as potential therapeutic agents due to their broad-spectrum activity and rapid, membrane-targeting mechanisms.[9–11] Multiple clinical efforts have explored their potential to combat bacterial infections.[11–14] Among these, the human cytolytic protein granulysin (GNLY), produced by cytotoxic T lymphocytes (CTLs) and natural killer (NK) cells, plays a key role in host defense against intracellular bacterial pathogens.[15] GNLY exhibits potent antimicrobial activity against Gram-negative, Gram-positive bacteria as well as certain fungi and parasites [16–19]. GNLY’s positively charged helical structure interacts directly with microbial cell membranes, enhancing permeability and causing lysis.[20, 21] This selectivity toward cholesterol-poor bacterial membranes reduces toxicity toward to mammalian cells.[22–24] Despite these advantages, the clinical translation of GNLY and related AMPs remains limited. Key challenges include susceptibility to proteolytic degradation,[25, 26] reduced activity under physiological salt conditions,[20] limited bioavailability,[13] and potential cytotoxicity at therapeutic doses.[27] These limitations highlight the need for delivery systems that can stabilize AMPs, enhance their activity under physiological conditions, and enable site-specific release.

Lipid-based delivery systems have emerged as promising platforms for improving the stability and efficacy of bioactive molecules.[28] In particular, self-assembled structures formed by amphiphilic lipids, including non-lamellar lyotropic liquid crystalline (LLC) phases, for tunable encapsulation and release.[29] Such systems can be engineered to respond to specific environmental stimuli, such as pH, salt concentration, or enzymatic activity, enabling controlled release at sites of infection.[28, 30–40] Clinically approved lipid formulations, such as AmBisome (liposomal amphotericin B), demonstrate the clinical potential of this approach.[41] However, challenges including instability under physiological conditions, immune clearance, and manufacturing complexity motivate the development of simpler and more robust systems.

This study demonstrates a pH-responsive lipid-protein self-assembly strategy based on OA and GNLY. OA was selected as a lipid component due to its amphipathic nature, biocompatibility, and ability to self-assemble with cationic proteins, forming stable nanostructures under physiological conditions.[42] We hypothesize that electrostatic and hydrophobic interactions between negatively charged OA and cationic GNLY enable the formation of co-assembled structures and neutral pH, while acidification triggers structural reorganization and release of GNLY. This concept is relevant for infected tissue where bacterial metabolism can lead to acidification.[43–45] The acidic microenvironment of infected tissues, such as acutely infected skin, provides an opportunity for pH-targeted release to improve therapeutic outcomes.[46] Similar ph_responsive lipid-LL37 systems have been demonstrated, supporting the feasibility of this approach.[47]

The self-assembly, structure, and pH-responsive behavior of OAGNLY nanocarriers are characterized using a combination of biophysical and biochemical techniques. The antimicrobial efficacy is evaluated against a range of clinically relevant MDR bacteria under physiological conditions, alongside assessments of cytocompatibility and therapeutic performance. In addition, the therapeutic potential of OAGNLY is investigated in a murine surgical wound-infection model, including analysis of reductions in bacterial load, histopathological changes, and biocompatibility. By integrating lipid self-assembly with innate immune effector function, this work aims to establish a platform for localized, infection-responsive antimicrobial therapy.

## 2 Results

### 2.1 Colloidal structure of OA and GNLY dispersions

Dispersed particles containing structures with different OA to GNLY weight ratios between 1:0 and 1:1 in their core were analyzed using SAXS at pH 7.0 and 5.0 (Fig. 1a and graphical illustration in supplementary Fig. 1a). The SAXS curve of OA alone (1:0) at pH 7.0 shows Bragg reflections at q-values characteristic of a dispersed micellar cubic phase with Fd3m symmetry. The corresponding lattice parameter, *a*, calculated from the 220 reflection at *q* = 1.30 nm^-1,^ yields *a* = 13.1 nm, in agreement with previous findings [42]. The addition of GNLY at GNLY/OA = 9:1 induced a shift of the peaks to slightly lower q-values, indicating a swelling of the Fd3m structure inside the dispersed droplets (Supplementary Fig.1b). Further increase to 7:3 and 1:1 induced a phase change. The SAXS pattern at 1:1 displays a Bragg reflection at q ≈ 0.80 with further, broader peaks at and above q = 1.35 nm⁻¹. The spacing is approximately consistent with hexagonal ordering, although the limited number of peaks precludes a definitive structural assignment. Nevertheless, the appearance of this distinct scattering pattern compared to the pure OA system demonstrates that GNLY interacts with and becomes incorporated into the lipid nanostructure, inducing a measurable structural rearrangement.

**Figure 1.**
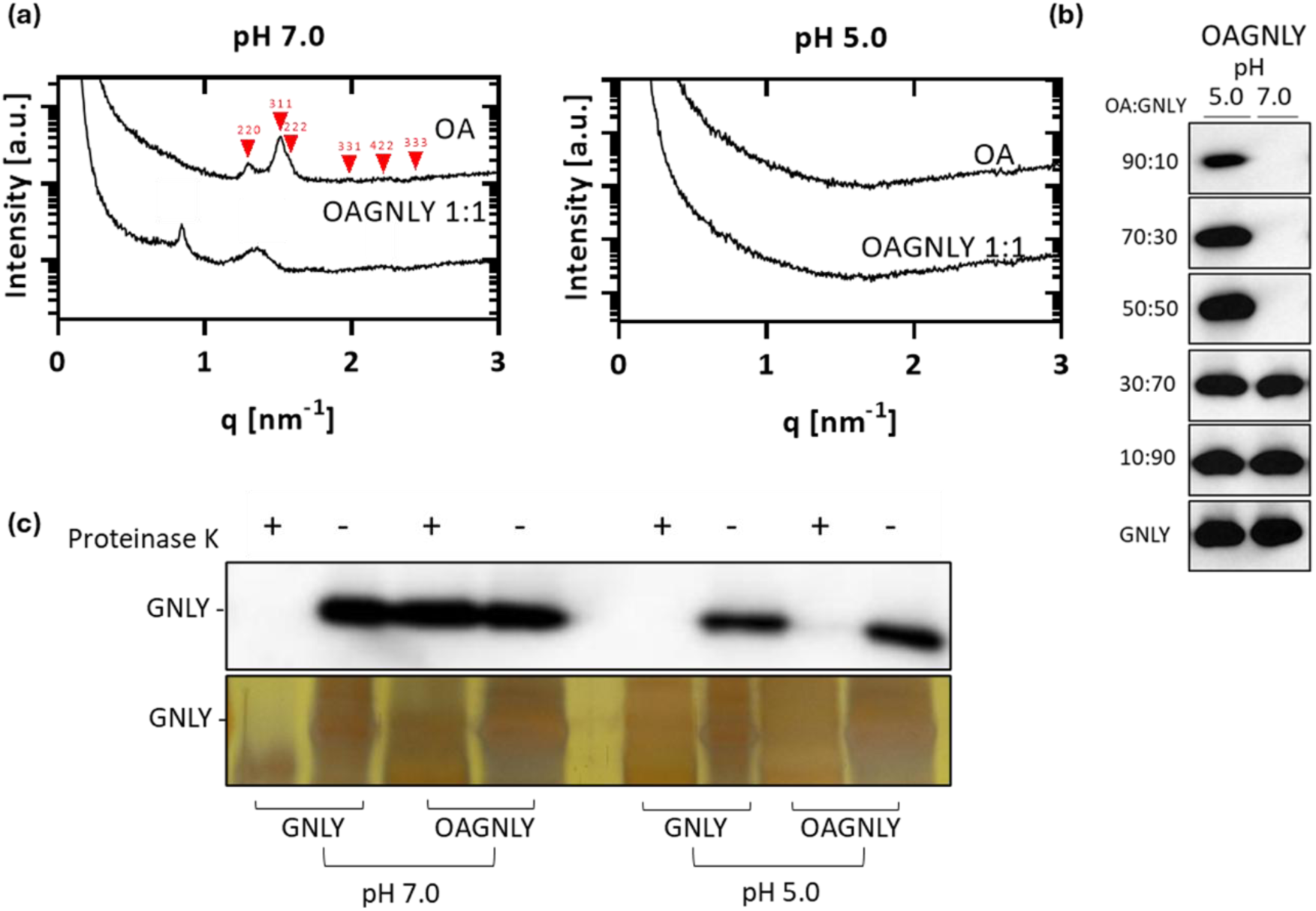
Characterization of OAGNLY self-assemblies. **(a)** SAXS data of OAGNLY nanocarriers at a 1:1 weight ratio (OA:GNLY) highlighting the structural transitions between pH 7.0 and 5.0. At pH 7.0, Bragg reflections corresponding to the Fd3m micellar cubic phase (red) were indexed with Miller indices, with a lattice parameter of 13.1 nm. Upon increasing the GNLY content, a phase transition was observed. At pH 5.0, the scattering curves are characteristic of unstructured emulsions. Representative data from three independent experiments are shown. For SAXS data of additional OAGNLY ratios, see Supplementary Fig. 1. **(b)** pH-triggered release of GNLY assessed by Western blot analysis after spin filtration (100 kDa Molecular weight cut off (MWCO)) of OAGNLY nanocarriers. At pH 7.0, GNLY was retained in the nanocarriers for all formulations tested (up to saturation at OA:GNLY ≥ 30:70). At pH 5.0, GNLY was detected in the filtrate across all formulations, indicating pH-dependent release. **(c)** Proteinase K treatment was used to assess the proteolytic stability of GNLY and OAGNLY at pH 7.0 and 5.0. The top panel shows a Western blot detecting GNLY, and the bottom panel represents the corresponding silver staining gel. GNLY alone exhibited significant degradation upon exposure to proteinase K, as evidenced by diminished bands. In contrast, GNLY encapsulated within OAGNLY at pH 7.0 was protected from proteolytic degradation, with intact bands observed. These results highlight the protective effect of OAGNLY in shielding GNLY from enzymatic degradation.

The SAXS profile of the OAGNLY system at pH 5.0 showed only a low-q upturn associated with scattering from emulsion droplets, similar to the one seen in OA in water emulsions at this pH value (Fig. 1a and Supplementary Fig. 1b).[42] The pH-dependent change in internal structure observed by SAXS shows a transition from internally nanostructured lipid particles containing GNLY and water at pH 7.0 to conventional emulsions lacking internal water and protein at pH 5.0. This structural transition suggests that acidification triggers the release of GNLY from the lipid particles.

The pH-dependent size distribution of OAGNLY particles at a 1:1 ratio was evaluated using multi-angle dynamic light scattering (DLS) (Supplementary Fig. 2a). At pH 7.0, the apparent hydrodynamic radius (R_H_), calculated across different scattering angles, ranged from 96 to 140 nm, with an average polydispersity index (PDI) of 0.24. In contrast, at pH 5.0 the R_H_ within the same angular range increased to 374–560 nm, with an average PDI of 0.12. The relatively low PDI values at both pH conditions indicate a largely monomodal particle size distribution. The larger particle size observed at pH 5.0 is consistent with the nanostructural transformations inferred from SAXS, supporting the hypothesis of a transition from internally nanostructured emulsion droplets to conventional emulsions. Notably, whereas OA particles aggregated within minutes at pH ≈ 5.0, the OAGNLY system remained colloidally stable for several days, further supporting a stabilizing role of GNLY. This stabilization likely arises from interactions between GNLY and oleic acid at the droplet interface, which prevent rapid coalescence under acidic conditions while still permitting the structural rearrangements associated with pH-triggered GNLY release.

Electrophoretic mobility measurements provided insights into GNLY integration into OA-based carriers (Supplementary Fig. 2c). At pH 7.0, OA particles exhibited a ζ-potential of −16 mV, consistent with stabilization by electrostatic repulsion among negatively charged, deprotonated OA molecules. Upon progressive addition of GNLY (OA:GNLY ratios from 9:1 to 1:1), the ζ-potential increased to −6 mV, indicating incorporation of the cationic GNLY into the assemblies. At pH 5.0, the ζ-potential became slightly positive (+0.46 mV).

The pH-dependent morphology of the OAGNLY nanocarriers was further examined using transmission electron microscopy (TEM) with negative staining. At pH 7.0, the nanocarriers formed smaller particles of approximately 50-150 nm (Supplementary Fig. 3a), consistent with DLS measurements when accounting for the larger hydrodynamic size obtained by DLS compared to dry-state TEM. In contrast, at pH 5.0, OAGNLY nanocarriers formed larger emulsion droplets of around 300–500 nm (Supplementary Fig. 3b). Together, these observations confirm the pH-dependent structural and morphological differences of the OAGNLY nanocarriers.

To further investigate the GNLY association with the nanocarriers and pH-triggered release, free GNLY in the dispersion was quantified using spin filtration (100 kDa MWCO) followed by Western blot analysis (Fig. 1b). At pH 7.0, GNLY was not detected in the filtrate for OA:GNLY ratios ranging from 9:1 to 1:1, indicating efficient retention of GNLY within the OA-based nanocarriers. At higher GNLY loadings (e.g., 3:7 and above), GNLY became detectable in the filtrate, suggesting saturation of the carrier system. In contrast, at pH 5.0, free GNLY was consistently detected in the filtrates across all formulations, consistent with pH-triggered release from the nanocarriers.

To assess whether encapsulation protects GNLY from proteolytic degradation, free GNLY or GNLY encapsulated in OAGNLY nanocarriers was treated with proteinase K, a broad-specificity serine protease. Free GNLY was readily degraded at pH 5.0 and 7.0. In contrast, GNLY encapsulated in OAGNLY was protected from proteolysis at pH 7.0 but not at pH 5.0, demonstrating the protective effect of encapsulation under neutral conditions (Fig. 1c). These findings are consistent with SAXS, DLS, and electrophoretic mobility measurements, confirming efficient encapsulation at pH 7.0 and pH-triggered release under acidic conditions.

### 2.2 OAGNLY carriers efficiently kill multidrug-resistant bacteria *in vitro* without triggering resistance development during prolonged treatment stress

The antibacterial efficacy of OAGNLY carriers was evaluated against a panel of multidrug-resistant bacteria at pH 5.0 and 7.0 by monitoring bacterial growth via turbidity measurements (OD600) (Fig. 2a-f). The colony forming units (CFUs) following treatment were estimated by comparison to reference growth curves generated from serial bacterial dilutions (Supplementary Fig. 4). Routinely, the findings of the growth assays were validated by spotting dilution assays (Supplementary Fig. 5). Given that GNLY activity is sensitive ionic strength and is completely inhibited at 150 mM NaCl [20], antimicrobial activity against MRSA and colistin resistant *E. coli* were conducted at both physiological (154 mM) and low (50 mM) NaCl concentrations.

**Figure 2:**
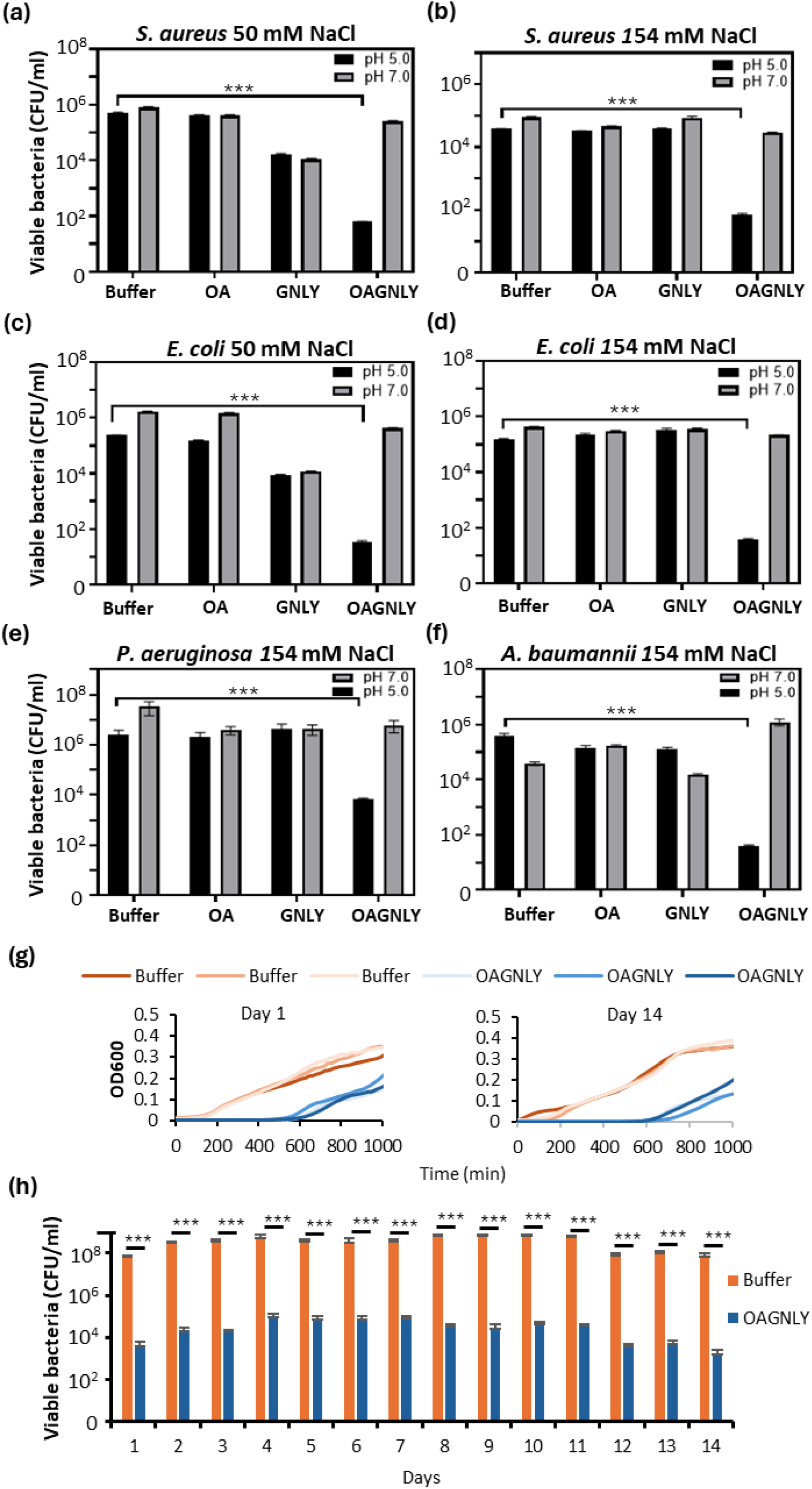
In vitro bactericidal activity of OAGNLY carriers. **(a-f)** Bactericidal activity of OAGNLY nanocarriers against methicillin resistant S. aureus (MRSA), colistin resistant E. coli (CREC) as well as carbapenem resistant P. aeruginosa and A. baumannii at varying pH (5.0 and 7.0) and salt concentrations (50 mM and 154 mM NaCl). Multidrug-resistant bacteria were treated for 2 hours with OAGNLY (32 µg/mL total = 16 µg/mL OA and 16 µg/mL GNLY), GNLY (32 µg/mL), OA (32 µg/mL), or PBS (control) at indicated pH and salt concentrations, followed by dilution in LB or BHI broth. Growth curves were monitored for 24 hours using a microplate reader, and CFUs were calculated from the growth curves (Supplementary Fig. 4). **(a)** MRSA at 50 mM NaCl, **(b)** MRSA at 154 mM NaCl, **(c)** CREC at 50 mM NaCl, **(d)** CREC at 154 mM NaCl, **(e)** carbapenem resistant P. aeruginosa, and **(f)** A. baumannii at 154 mM NaCl. MRSA were repeatedly treated for 14 days with OAGNLY (32 µg/mL) before viability assessment in growth assays. Representative growth curves (in triplicates) at day 1 and day 14 are presented in **(g)** and the quantification of the results are shown in **(h)**. Bars represent CFU means ± SEM on a log scale from three independent experiments, each performed in triplicate. Statistical significance was determined by ANOVA: *P < 0.05, **P < 0.01, ***P < 0.005.

At 50 mM NaCl (Fig. 2a, c), free GNLY (32 µg/mL) reduced the viable bacterial population by approximately 1 log at pH 5.0 and 7.0, while 32 µg/mL OAGNLY (16 µg/mL OA, 16 µg/mL GNLY) carriers achieved a more than 3-log reduction at pH 5.0. At 154 mM NaCl (Fig. 2b, d), only OAGNLY carriers (32 µg/mL) maintained significant bactericidal activity against MRSA and CREC at pH 5.0. Neither GNLY (32 µg/mL) nor OA (32 µg/mL) alone affected bacterial viability at physiological salt concentrations. Notably, OAGNLY (32 µg/mL) carriers showed no activity at pH 7.0 under any tested conditions. Similarly, OA alone did not significantly impact bacterial growth across all tested concentrations (0.01–10 mg/mL) at either pH.

The bactericidal activity of OAGNLY (32 µg/mL) was further assessed against carbapenem resistant *Pseudomonas aeruginosa* and *Acinetobacter baumannii* under physiological salt concentration (154 mM NaCl) (Fig. 2e, f). After 1 hour of treatment at pH 5.0, OAGNLY (32 µg/mL) achieved a 3-log reduction against *Pseudomonas aeruginosa* (Fig. 2e) and 4-log reduction in *Acinetobacter baumannii* (Fig. 2f). Together, these results demonstrated that OAGNLY nanocarriers restore antimicrobial activity under physiological salt conditions and enable potent bactericidal effects against a range of clinically relevant MDR pathogens.

Minimal inhibitory concentrations (MICs) of OAGNLY carriers under physiological salt conditions were determined to be 16 µg/mL for MRSA and 32 µg/mL for colistin-resistant *E. coli* at pH 5.0 (Supplementary Fig. 6). No inhibitory activity was observed for GNLY or OA alone under these conditions.

Antimicrobial activity against colistin-resistant *E. coli* was further assessed using a microbial cell viability assay that measures ATP production reflecting metabolic activity (Supplementary Fig. 7). Luminescence, which correlates to cellular ATP content, was significantly reduced in samples treated with OAGNLY (32 µg/mL) at pH 5.0 compared to those treated with OA (32 µg/mL), GNLY (32 µg/mL), or PBS controls. This indicates a substantial reduction in bacterial viability when exposed to OAGNLY, confirming the growth analyses.

An important question was if continuing sublytic treatment stress might lead to the development of resistance against OAGNLY nanocarriers. Therefore, to ensure sufficient surviving bacteria after every treatment cycle, we treated a rather high bacterial concentration (10^7^ to 10^8^ MRSA/ml) with 32 μg/ml OAGNLY in physiological salt at pH 5.0 for 2 hours before diluting in broth to monitor growth for 24 hours (Fig. 2g). The surviving bacteria were repeatedly treated for two weeks. In these two weeks of continued survival stress in this population, the bactericidal activity of OAGNLY did not change, indicating no detectable emergence of resistance under the tested conditions (Fig. 2h)

Transmission electron microscopy (TEM) revealed pronounced ultrastructural damage in *E. coli* following exposure to OAGNLY nanocarriers (32 µg/mL) in PBS at pH 5.0, including disruption of the cell envelope and cytoplasmic structures and the nucleoid structure (Fig. 3 and Supplementary Fig. 8a). These features were not observed in control-treated samples. At pH 7.0, OAGNLY particles were predominantly observed adhering to the bacterial surface (red arrows, Fig. 3b and Supplementary Fig. 8b). Larger structures (100–400 nm) at pH 5.0 likely correspond to OA-rich emulsion droplets following GNLY release and/or remnants of lysed bacterial cells.

**Figure 3:**
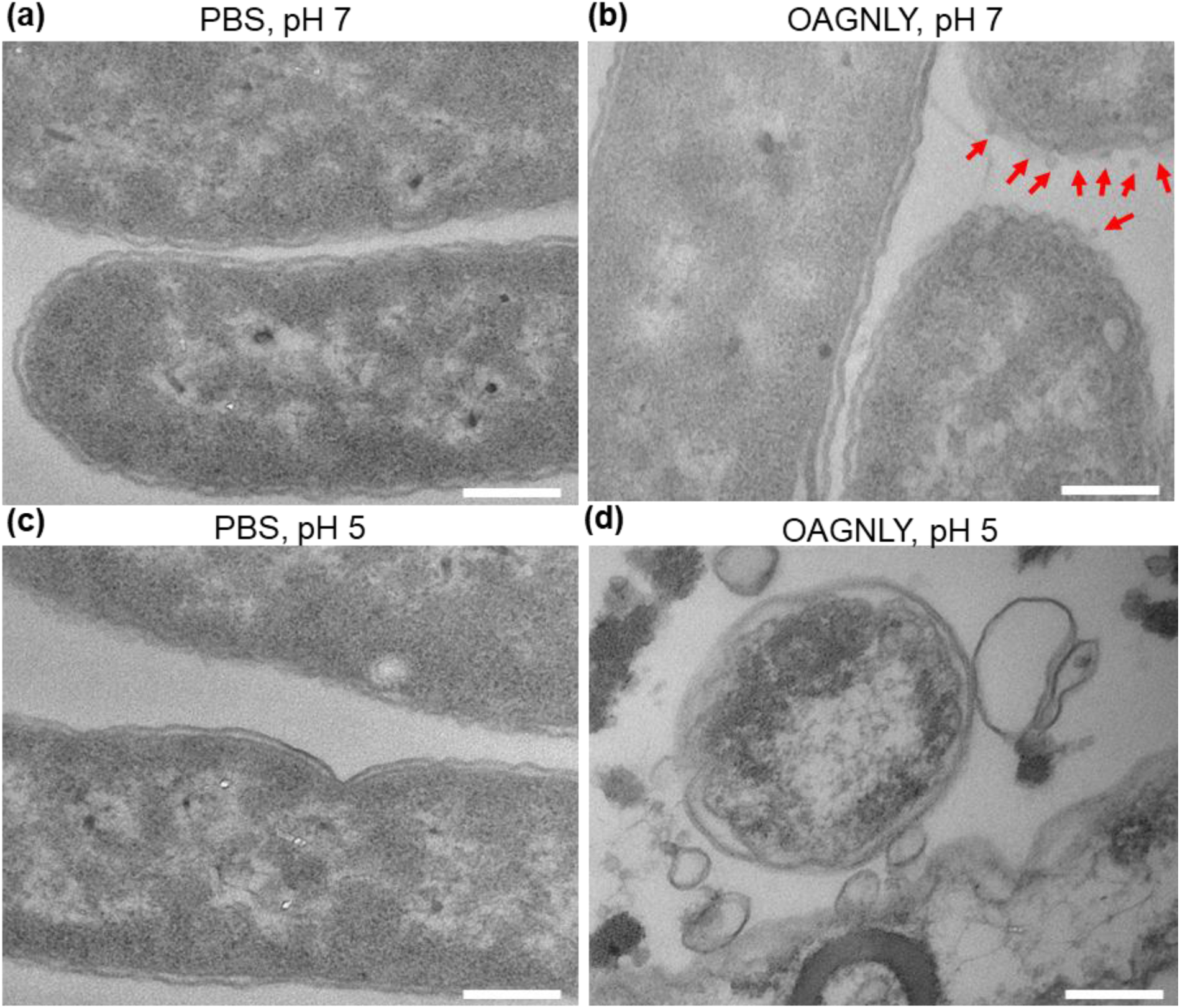
Morphological alterations of E. coli upon treatment with OAGNLY carriers. **(a, c)** Untreated E. coli cells in PBS containing 154 mM NaCl at pH 7.0 and pH 5.0, respectively, exhibit intact membranes and cytoplasmic structures. **(b, d)** E. coli treated with OAGNLY carriers (32 µg/mL) show significant ultrastructural changes. At pH 7.0 **(b)**, red arrows highlight the attachment of OAGNLY nanocarriers to the outer bacterial membrane. At pH 5.0 **(d)**, profound damage to the bacterial membrane and cytoplasmic contents was observed, indicating severe disruption and cell death. Representative high-resolution TEM images are provided from three independent experiments. Bars = 200 nm. Lower-resolution images are available in Supplementary Fig. 7.

To further investigate the interaction of OAGNLY nanocarriers with bacterial surfaces, high resolution confocal microscopy was performed using GNLY tagged with a fluorescent dye (Alexa Fluor 488) and incorporated into OAGNLY nanocarriers. In *E. coli* treated with these fluorescently labeled OAGNLY nanocarriers, strong fluorescence signals were observed at the bacterial surface, indicating association of the nanocarriers with the bacterial envelope (Supplementary Fig. 8a). Higher magnification revealed heterogeneous binding patterns with some cells displaying clustered fluorescence signals and others showing discrete dots (arrows in Supplementary Fig. 8b). In contrast, fluorescently labeled GNLY alone displayed minimal attachment under physiological salt concentration consistent with weaker interaction in the absence of OA (Supplementary Fig. 8c). These findings support enhanced interactions of OAGNLY nanocarriers with bacterial surfaces and are consistent with a mechanism involving pH-triggreed GNLY release and membrane disruption.

### 2.3 OAGNLY nanocarriers display cytocompatibility and reduced bacterial burden *in vivo*

The cytocompatibility of OAGNLY carriers was first evaluated *in vitro* using an MTS (3-(4,5-dimethylthiazol-2-yl)-5-(3-carboxymethoxyphenyl)-2-(4-sulfophenyl)-2H-tetrazolium) assay to assess metabolic activity and viability in HeLa cells. Cells were treated with OAGNLY 32 µg/mL (16 µg/mL OA and 16 µg/mL GNLY) for 2 and 4 hours at pH 5.0 and 7.0, respectively (Supplementary Fig. 10). No additional reduction in cell viability was observed compared to PBS-treated controls. However, prolonged incubation at pH 5.0 alone reduced cell viability, limiting the duration of cytotoxicity assessments under acidic conditions.

To further evaluate cytocompatibility under physiological conditions, human primary dermal fibroblasts (HDF) and HeLa cells were treated for 24 hours at pH 7.0. At the bactericidal concentration of 32 µg/mL (16 µg/mL OA and 16 µg/mL GNLY), neither OAGNLY carriers nor its individual components induced significant cytotoxicity compared to the pro-apoptotic positive control, staurosporine (STS) (Fig. 4a). These results indicate that OAGNLY formulations are well tolerated by mammalian cells at concentrations effective for bacterial killing.

**Figure 4:**
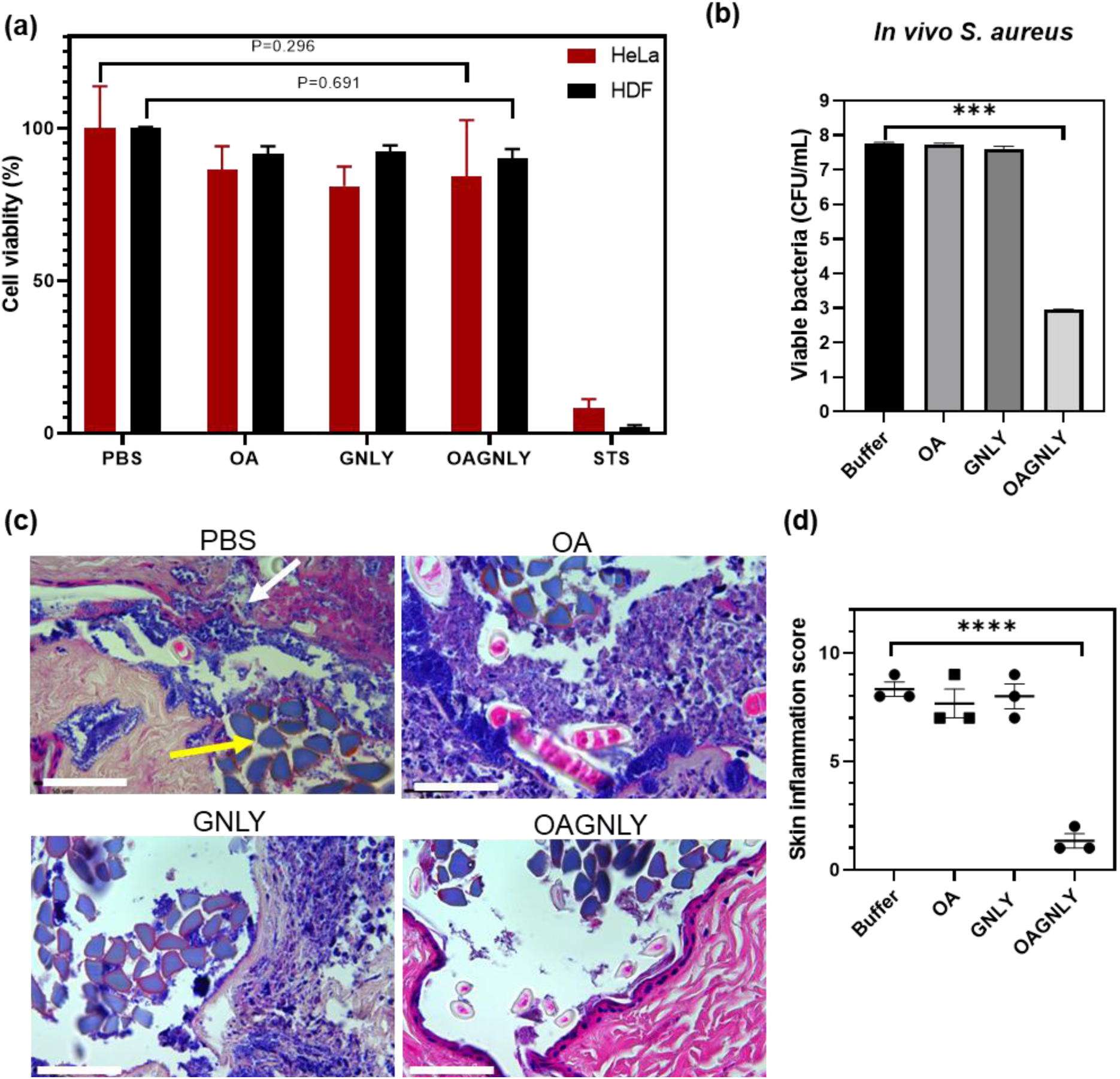
Toxicity against mammalian cells and in vivo studies. **(a)** Cytotoxicity of OAGNLY and its individual components was assessed using HDF and HeLa cells at pH 7.0 after 24 hours of treatment. Cell viability after indicated treaements (32 μg/ml) was determined via MTS assay, and values were normalized to PBS-treated controls. Bars represent the mean percentage of cell viability ± SEM from three independent experiments. The results indicate that OAGNLY nanocarriers display high biocompatibility, demonstrating their safety for potential therapeutic applications. **(b)** Balb/c mice were infected with MRSA contaminated sutures in surgical sites for 1 hour before treatment with OAGNLY (32 µg/mL) or individual components (OA = 32 µg/mL and GNLY = 32 µg/mL) for 4 hours. MRSA bacterial load in excised and homogenized surgical site wounds was quantified by CFU spot dilution assays. Bars display mean CFUs ± SEM based on three independent experiments (n = 5 mice per group). The significant reduction in CFUs in the OAGNLY-treated group underscores its potent activity against multidrug-resistant bacteria in a surgical site infection model. **(c)** Skin biopsies from treated mice were analyzed for bacterial load, tissue penetration, inflammation, and vascular changes using H&E and Gram staining. Representative images from three independent experiments are shown. In the PBS-treated group, the white arrow indicates bacterial invasion, while the yellow arrow highlights the suture in the tissue. **(d)** Inflammation scores were determined in a blinded manner by evaluating parameters such as skin thickness, abscess formation, bacterial penetration depth, immune cell infiltration, and changes in the microvasculature and epidermis across three independent experiments (n = 5 mice per group). The graph shows dotted means for individual experiments alongside the average ± SEM. Refer to supplementary Figures 10 and 11 for additional data. Statistical significance was calculated using ANOVA, with *P < 0.05, **P < 0.01, ***P < 0.001, ****P < 0.0001.

The antimicrobial efficacy of OAGNLY was next assessed in a surgical skin infection model. MRSA-infected sutures were used to establish localized bacterial infections for 1 hour prior to topical treatment with OAGNLY nanocarriers (32 µg/mL), OA, GNLY, or PBS for 4 hours. Bacterial load was quantified by CFU analysis of homogenized skin tissue samples. OAGNLY treatment resulted in > 5 log reductions in bacterial burden, approaching the detection limit (Fig. 4b), whereas OA or GNLY alone showed no significant antimicrobial activity. These findings were confirmed by growth curve analyses of recovered bacteria (Supplementary Fig. 11).

Histopathological analysis of treated skin revealed markedly reduced inflammation and bacterial penetration in OAGNLY-treated tissue compared to control groups (Fig. 4c and Supplementary Fig. 12 and 13). In contrast, PBS, OA, or GNLY treated samples exhibited widespread bacterial presence in the epidermal and deeper tissue layers, accompanied by pronounced inflammatory responses, including edema and erythema. In contrast, OAGNLY-treated skin displayed minimal signs of inflammation and negligible bacterial presence.

Quantitative assessment of inflammation showed that skin thickness increased from ∼540 µm in non-infected mice to 1000–1400 µm in infected controls. OAGNLY-treated mice exhibited reduced skin thickness values of 560–800 µm (Supplementary Fig. 12). In addition, inflammation was scored using criteria such as abscess formation, inflammatory cell infiltration, bacterial penetration depth, and vascular alterations[48] (Supplementary Fig. 12 and 13). OAGNLY treated samples showed significantly reduced inflammation scores compared to controls (Fig. 4d). These results demonstrate that OAGNLY nanocarriers effectively reduce bacterial burden and associated inflammation *in vivo*, while maintaining favorable cytocompatibility, supporting their potential as a platform for localized antimicrobial therapy.

## 3 Discussion and conclusion

The increasing prevalence of AMR represents a major global health challenge and underscores the urgent need for innovative therapeutic strategies.[3] This study demonstrates that OA and GNLY can form pH-resposive self-assembled nanocarriers that enable retention of GNLY under neutral conditions and its release under acidic environments relevant to infection sites. This behavior provides a mechanism for localized activation of the antimicrobial system.

SAXS and ζ-potential measurements indicated that GNLY association with OA is primarily driven by electrostatic interactions between the negatively charged carboxyl groups of OA and the positively charged (arginine) residues of GNLY.[49] The observed shift in ζ-potential from −16 mV to −6 mV upon GNLY addition supports this interaction. At neutral pH around 7.0, partial deprotonation of OA [42] promotes structured assemblies capable of retaining GNLY, whereas at acidic pH around 5.0, protonation of OA disrupts these structures, leading to the formation of unstructured emulsions and release of GNLY. This pH-dependent transition is consistent with SAXS and release data and provides a mechanism for infection-responsive delivery. While GNLY retention at pH 7.0 and increased detection at pH 5.0 support this model, the exact localization of GNLY within the assemblies remains to be further elucidated. The acidic microenvironments of bacterial infections might facilitate the targeted release of GNLY at the site of infection.[36, 46]

Functionally OAGNLY nanocarriers restored GNLY activity under physiological salt conditions, where the free GNLY is largely inactive. Enhanced antimicrobial activity at pH 5.0 and inactivity at pH 7.0 may result from a combination of GNLY release and increased electrostatic interactions between carriers and negatively charged bacterial membranes.[16, 20] The observed activity across MDR pathogens, including MRSA, *Pseudomonas aeruginosa*, and *Acinetobacter baumannii*, highlights the broad applicability of this approach. Repeated sublethal exposure over two weeks did not result in detectable changes in susceptibility under the conditions tested, suggesting a low propensity for resistance development, although longer-term studies will be required to confirm this. This was recently also shown for prolonged GNLY treatments in combination with granzyme B. [50]

In contrast to certain last-resort antibiotics, such as colistin, which are associated with significant cytotoxicity,[6] OAGNLY nanocarriers exhibited favorable cytocompatibility in vitro at bactericidal concentrations. Furthermore, in a murine surgical wound infection model, OAGNLY treatment resulted in substantial reductions in bacterial burden and inflammation compared to controls. These findings support the potential of OAGNLY as a locally applied antimicrobial strategy for skin and soft tissue infections.

Despite these promising results, several limitations should be considered. The release behavior of GNLY was assessed qualitatively, and a more detailed quantitative analysis of loading efficiency and release kinetics would further strengthen the mechanistic understanding. In addition, while cytocompatibility was demonstrated *in vitro*, a comprehensive safety evaluation, including long-term and repeated-dose studies, will be required for clinical translation. Finally, direct comparison with standard-of-care antimicrobial treatments would help further contextualize the therapeutic performance of this system.

In summary, this study demonstrates that lipid-protein self-assembly between OA and GNLY enables the formation of pH-responsive nanocarriers that protect GNLY under neutral conditions and release it under infection-relevant acidic conditions. This strategy enhances GNLY stability and restores antimicrobial activity under physiological salt conditions, resulting in effective killing of multidrug-resistant bacteria and improved outcomes in a wound infection model. These findings establish a promising platform for infection-responsive antimicrobial delivery and highlight the potential of lipid-based self-assembly to improve the therapeutic performance of innate immune effectors.

## 4 Experimental Section

### Expression and purification of GNLY

Recombinant His-tagged GNLY was expressed and purified as described previously.[51] Briefly, GNLY was secreted from HEK293T cells and purified from the supernatant using nickel affinity chromatography (ÄKTA Prime; Amersham Biosciences, Uppsala, Sweden) with Ni-NTA (nitrilotriacetic acid) agarose resin (Qiagen, Hilden, Germany). The protein was eluted with 250 mM NaCl and 50 mM Tris using a linear imidazole gradient (0–1000 mM) at pH 8.0. Further purification was performed using a Mono S column (Cytiva #17516801), with elution via a linear NaCl gradient (0–1000 mM) in 50 mM HEPES, pH 7.4. GNLY purity was confirmed by SDS-PAGE followed by silver staining, and the presence of the His-tag was verified using a 6x-His tag antibody (HIS.H8) (Invitrogen #MA1-21315). For specific experiments, the His-tag was removed using enterokinase before S-column purification to ensure it did not interfere with ultrastructural studies. As an alternative, native GNLY was purified from YT Indy cells as described by Jerome Thiery *et al*.[52] The activity of every purified GNLY batch was routinely tested according to León, D. L *et al.* [53]

### Carrier Preparation

Lipid carriers were prepared by dispersing oleic acid (OA, 10 mg/mL; 99% purity, Sigma-Aldrich, Buchs, Switzerland) in phosphate buffer containing defined NaCl concentrations (pH 7.4). The dispersion was homogenized on a Lab500 NexTgen Ultrasonic platform (SinapTec, Lezennes, France) for 2 minutes in pulse mode (1 s on/1 s off) at 27% of the maximum amplitude (max power 500 W). Phosphate-buffered saline (PBS, 1×, pH 7.4) contained 137 mM NaCl (Acros Organics, 99.5% purity, Denmark). The pH was adjusted to 7.4 using freshly prepared 1 M NaOH or 1 M HCl solutions, made from NaOH pellets (>99% purity, Sigma-Aldrich) and concentrated HCl stock (∼37%, analytical grade, Fisher Scientific, USA). For granulysin (GNLY) lipid carrier formulations, GNLY was incorporated into OA dispersions at mass ratios of 1:1, 3:7, and 9:1. The pH was then adjusted to either 5.0 or 7.0, and samples were equilibrated at room temperature for 1 hour prior to analysis.

### Small angle X-ray scattering (SAXS)

Structural characterization of the carriers was conducted via SAXS at the Austrian SAXS beamline, ELETTRA Synchrotron, Trieste, Italy. Samples were prepared under different pH conditions with OA and GNLY at 1:1, 3:7, 9:1 ratios. Dispersions were transferred to thin-walled quartz capillaries, sealed, and placed in the sample holder. The samples were exposed to X-rays of wavelength (λ) 0.77 Å (16 keV), with a sample-to-detector distance of 1830 mm, providing a q-range of 0.1 < q < 7 nm⁻¹, where q is the scattering vector defined as is defined as 𝑞 = (4π/λ) sin(θ/2), where θ is the scattering angle.

### Multi-angle Dynamic light scattering (DLS)

DLS measurements were performed using a light-scattering goniometer (CGS-8F, ALV, Langen, Germany) and a solid-state laser (Coherent Verdi V5, 532 nm wavelength, max. power 5W). Data were collected with fiber-optic detection optics (OZ, GMP, Zürich, Switzerland), eight fiber-optic detectors, and ALV 7004 correlators with fast expansion (ALV, Langen, Germany). Scattering angles ranged from 39° to 124°. The apparent hydrodynamic radius (RH) was calculated from the apparent diffusion coefficient (*D_app_*) obtained by cumulant analysis of the DLS autocorrelation function using the Stokes-Einstein equation.[54] The polydispersity index (PDI) was calculated from the second cumulant. The average decay constant (Γ̅) was plotted against 𝑞^2^, and the slope of the resulting linear fit was used to determine the apparent translational diffusion coefficient (D).

### Electrophoretic Mobility measurements

The ζ-potential of the carriers was measured at pH 5.0 and 7.0 using phase analysis light scattering (PALS)[55] on a DelsaMax Pro Nano instrument (Beckman Coulter, Inc., Brea, USA). OA and GNLY dispersions at 0%, 10%, 30%, and 50% weight fractions were analyzed in 45 μL quartz cuvettes at 25°C. The ζ-potential was calculated using the Smoluchowski equation:

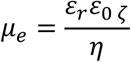

where *μ_e_* is electrophoretic mobility, *ε_r_* is the dielectric constant of water*, ε*_0_ the permittivity of vacuum and *η is the* viscosity of water.

### pH triggered release by fractionation and Western blot

OAGNLY samples at pH 5.0 and 7.0 were fractionated at 4000 xg at 25°C using 100 kDa centrifugal filters (Amicon Ultra-15). The retained and filtrated fractions were separated on 15% SDS polyacrylamide gels following standard procedures. Proteins were transferred onto polyvinylidene difluoride (PVDF) membranes using a Trans-Blot Turbo system (Bio-Rad). Membranes were blocked in TBS-Tween containing 1% bovine serum albumin (BSA) for 1 hour before incubation with the primary antibody (6x-His tag antibody, Invitrogen #MA1-21315) at 4°C overnight. After washing, membranes were incubated with a secondary antibody (anti-mouse IgG, RD Systems #HAF007) at 1:2000 dilution in a blocking buffer (3% BSA in TBS-Tween). Visualization was performed using a chemiluminescent substrate (Westar Sun #HA20A-MZ) and imaged using a gel documentation system (GBox, Syngene).

### *In vitro* antibacterial assays

Glycerol stocks of colistin-resistant *E. coli* strains (encoding MCR-1: 10 TC2, 23-1 TC, 57 TC3, G2 TC2) and a recipient non-resistant strain (J53), as well as methicillin-resistant *S. aureus* (MRSA, USA300), *Pseudomonas aeruginosa* (PAO1 and PA14) and *Acinetobacter baumannii* (ATCC19606) were streaked on plate count (PC) agar (Sigma-Aldrich, Buchs, Switzerland) and incubated overnight. Single colonies were cultured in Lysogeny broth (LB) broth (*E. coli*) or brain-heart infusion (BHI) medium (*S. aureus*) or Mueller Hinton broth (#70192 Millipore) overnight. Respective cultures were diluted 1:50 in fresh broth and grown to mid-log phase (eg. *E. coli* OD600 ≈ 0.4, ∼10⁹ CFU/mL). Bacteria were washed twice in sodium dihydrogen phosphate buffer and resuspended to ∼10^5^ to 10^6^ CFU/mL in assay buffer. For antibacterial testing, OAGNLY dispersions were prepared in 10 μL assay buffer in a transparent, flat-bottomed 96-well plate (Corning). Controls included buffer-only, 1-256 μg/mL GNLY or OA at pH 5.0 or 7.0. Each well received 10 μL of bacterial suspension (∼2.5 × 10⁵ CFU) and was incubated at 37°C for 1 hour. After treatment, 150 μL of broth was added, and bacterial growth was monitored in a plate reader (BioTek Synergy H1) with intermittent shaking (15-second intervals every 15 minutes) for 24 hours at OD600 nm. Growth curves were calibrated against reference curves obtained from 10-fold serial dilutions of bacteria grown for 24 hours. Simultaneous colony-forming unit (CFU) assays were conducted to determine the exact initial bacterial counts. Lag-phase times (defined as the time to reach an OD threshold of 0.1) were correlated with starting CFU to calculate bacterial numbers post-incubation.

### Determination of minimal inhibitory concentration (MIC) and the development of resistance

MIC values for GNLY, OAGNLY, and OA were determined by exposing bacteria (∼10^5^/ml) to two-fold serial dilutions (1–256 μg/mL) in 96-well plates. After 1 hour of incubation, 250 μL of Mueller Hinton broth (#70192 Millipore) was added to each well. Growth curves were recorded as described above. MIC was defined as the lowest concentration that inhibited bacterial growth completely after 24 hours, as determined by OD600 readings and CFU counts.[56]

For the determination of resistance development, 10^7^ to 10^8^/ml exponential phase MRSA were treated with 32 μg/ml OAGNLY in phosphate buffered saline (50 mM, adjusted to pH 5) for 2 hours before 10fold dilution in Mueller Hinton broth to monitor growth for 24 hours. The next day, using the regrown population of MRSA, the treatment procedure was repeated. The procedure was repeated 14 times.

### Proteolytic degradation

Proteinase K (20 µg/mL, Boehringer Mannheim GmbH) digestion was performed at pH 7.0 and 5.0 for 30 minutes at 37°C. Enzymatic activity was halted by heat inactivation. The presence or absence of OA and GNLY was tested in the samples. Resultant samples were separated using 15% SDS-polyacrylamide gel electrophoresis and transferred onto PVDF membranes. Western blotting was performed as described previously, using 6x-His tag antibody (Invitrogen #MA1-21315) as the primary and anti-mouse IgG (RD Systems #HAF007) as the secondary antibody.

### Preparation of Fluorescently Tagged GNLY and Fluorescence Microscopy

Granulysin (GNLY) was fluorescently labeled using a commercially available amine-reactive fluorescent dye (Alexa Fluor 488). Briefly, GNLY was incubated with the dye in a 1:1 molar ratio in 100 mM sodium bicarbonate buffer (pH 8.3) for 1 hour at room temperature. Unbound dye was removed using a desalting column, and the labeled GNLY was stored at 4°C in PBS. OAGNLY nanocarriers (GNLY and OA mixed in a 1:1 molar ratio, with GNLY fluorescently tagged). Each treatment was performed in PBS at pH 7.0 for 30 minutes at 37°C with gentle shaking. After treatment, the bacterial cells were washed three times with PBS to remove unbound compounds. The treated bacterial cells were fixed with 4% paraformaldehyde for 15 minutes at room temperature, washed with PBS, and mounted on glass slides using ProLong™ antifade mounting medium. Imaging was performed using a fluorescence microscope equipped with a 488 nm laser and an appropriate filter set. Images were acquired using a 63× oil immersion objective on Leica SP5 confocal microscope.

### Transmission electron microscopy (TEM)

*E. coli* treated with OAGNLY in physiological salt concentration was prepared as per antibacterial assay protocols. After treatment, bacteria were pelleted by centrifugation at 1200 × g for 5 minutes at 4°C. The pellet was fixed in 2.5% glutaraldehyde in PBS (pH 7.4) and washed with 0.1 M cacodylate sodium buffer (pH 7.4). Samples were embedded in 2% agar, centrifuged at 1200 × g, and trimmed for post-fixation in 1% osmium tetroxide. Dehydration was performed in a graded alcohol series followed by embedding in epoxy resin. Polymerization occurred at 60°C for 48 hours. Ultrathin sections (50 nm) were contrasted with lead nitrate and uranyl acetate. Samples were observed using a Philips CM100 Biotwin TEM (100 kV) equipped with a TVIPS F416 camera (4k × 4k) and SIS Morada camera.

### Negative staining TEM

Formvar/carbon-coated TEM grids were glow-discharged immediately before use to enhance hydrophilicity. OAGNLY samples at pH 5.0 and 7.0 were applied to the grid, which was held with reverse forceps, and incubated for 7 minutes. Excess sample was removed by blotting the grid at a 45-degree angle using filter paper (Whatman No. 1). The grid was then stained by incubating it for 7 minutes in 2% uranyl acetate, prepared on a Parafilm surface. Excess stain was similarly removed by blotting, and the grid was allowed to air dry for at least 30 minutes. Dried grids were stored in a grid box at room temperature in a dark environment until imaging. The prepared grids were then imaged using a Philips CM100 Biotwin TEM (100 kV) under standard conditions.

### Cell viability assay

Cell viability was measured using an MTS assay (Abcam #ab197010) with human dermal fibroblasts (HDF, Cell Applications #106K-05a) and HeLa cells. HDF cells were seeded in 96-well plates (5 × 10³ cells/well in 100 µL Fibroblast Growth Medium, Cell Applications #116-500), and HeLa cells in DMEM (Pan Biotech, P04-04510), supplemented with 10% heat-inactivated FBS (Sigma) and 1% antibiotic/antimycotic solution (Thermo Fisher). Cells were treated with OAGNLY and controls (OA, GNLY, or PBS) at indidcated concentrations and timees. Staurosporine (1 μg/ml, STS, AM-2282) served as the positive control. MTS reagent was added, and viability was measured at 490 nm using a microplate photometer (BioTek Synergy H1) after 15 minutes of incubation.

### In vivo studies

#### Mouse Model of Surgical Site Infections (SSIs)

*Preparation of Infected Sutures*: *Staphylococcus aureus* (MRSA) was cultured in 3.7% BHI broth at 37°C, 250 rpm overnight. Bacteria were pelleted, resuspended, and diluted to 2 × 10⁹ CFU/mL. Silk sutures (Assut #REF 643F) were soaked in the bacterial suspension for 30 minutes, dried, and stored at 4°C.[57]

*Infection Procedure*: All animal experiments were performed after the approval of the local Ethics Committee for Animal Studies (Approval number: 2022-16-FR). The animals were kept in a 12-hours light–dark cycle with free access to water and pellets (Granovit AG #0000111152).

BALB/c mice (25–30 g) were anesthetized with isoflurane (PerkinElmer #800-762-4000) and given buprenorphine (0.1 mg/kg) pre-surgery for pain relief. Two 1 cm incisions were made on each flank of the back, and 1 cm infected sutures were secured in the wounds. After 1 hour to establish a skin and soft-tissue infection, 30 µL OAGNLY, OA, GNLY (all in PBS), or PBS alone was applied for 4 hours. All mice were euthanized 5 hours post-infection. The pH of the GNLY/OA was kept at 7.0 before application. No deaths were reported in any infected mice.

*Sample Collection and CFU Enumeration*: A 2 × 1 cm area around the wound was excised, homogenized, and centrifuged. The pellet was washed and resuspended in PBS (pH 7.0). Diluted suspensions were plated on BHI agar (containing streptomycin, 50 µg/mL) and incubated at 37°C overnight for CFU determination. Growth curves were simultaneously recorded.

#### Histology and immunohistochemistry

Formalin-fixed, paraffin-embedded 1 µm skin sections were analyzed by hematoxylin and eosin (H&E) staining and bacterial Gram staining.[58] A histological scoring system (adapted from Rigoni et al.)[48] evaluated skin parameters, including dermal thickness, abscesses, immune cell infiltration, bacterial penetration, and epidermal alterations, scored on a scale of 0–3. Dermal thickness was measured using NanoZoomer (NDP view 2.3.10) and Leica Application Suite (LASX). The double-blind study used the following parameters: 1. Skin thickness (increases with the grade of inflammation), 2. Cutaneous abscesses, 3. Increased immune cells in dilated microvasculature and infiltrated in the dermis, 4. Bacterial penetration, and 5. Alteration of the epidermis. Scoring from 0 to 3 (where 0 is normal condition and 3 is severe structural changes) was established to evaluate different skin manifestations. Histological dermal thickness was determined, on H&E-stained sections, by measuring the distance between the epidermal-dermal junction and dermal–subcutaneous fat junction.

### Statistical analyses

For cell viability assays, two-way ANOVA was used for comparisons. For *in vitro* and *in vivo* data, one-way ANOVA was performed. Results are expressed as mean ± SD from triplicates (*in vitro)* from biological replicates. Six mice per group were used for in vivo assessment. Separate scientists were conducting experiments and analysing data in a blinded manner. Significance levels are denoted as follows: *p < 0.05, **p < 0.01, ***p < 0.005.

## Data availability

Data acquired and analyzed during this study are included in the manuscript, and supplementary information is available upon request from the corresponding authors.

## Acknowledgements

We thank Brigitte Scolari for her technical assistance with the transmission electron microscopy sample preparations and Marlène Sanchez for her technical assistance with the histological skin sections. Multidrug resistant bacteria strains were a kind gift of Dr. Laurent Poirel and Prof. Patrice Nordmann at the University of Fribourg and of the associated National Reference Center for Antibiotic Resistance (NARA). Research reported in this publication was supported by the SNSF through the NCCR Bioinspired Materials and grants IZBRZ2_186251, 200021_192051 to SS, and the Swiss National Science Foundation (SNSF grant # 31003A_182729), the Novartis Foundation for Medical-Biological Research and the Vontobel-Foundation to MW.

## Conflict of Interest

The authors declare no conflict of interest.

## Author contributions

Project conceptualization and study supervision: MW, SS. Design and ultrastructural analysis of carriers: OAH, MG. *In vitro* antimicrobial analysis: OAH, JV. Protein synthesis: OAH, PM. *In vivo* antimicrobial and histopathology analysis: OAH, OC. Manuscript writing: OAH, MW and SS.

## Supporting information

**Supporting figure 1.**
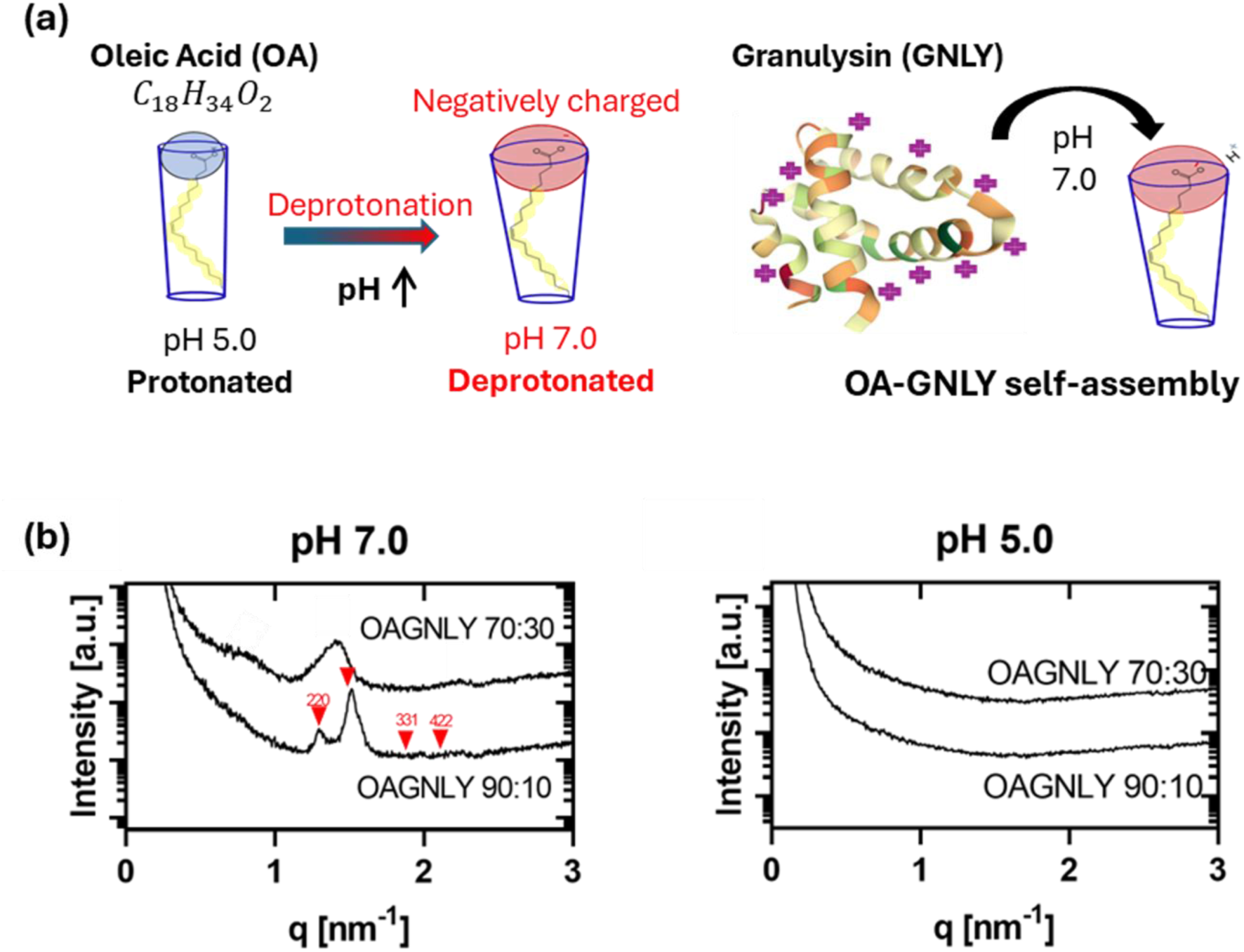
Graphical illustration of self-assembly formation and SAXS analyses at different OA to GNLY ratios. **(a)** OA is negatively charged due to deprotonation at pH 7 and therefore self-assembles with the positively charged GNLY. SAXS curves are shown for OAGNLY nanocarriers at concentration ratios of OA to GNLY of 90:10 and 70:30 at **(b)** pH 7.0 and **(c)** pH 5.0. At pH 7.0, distinct Bragg reflections corresponding to highly ordered structures are observed. These reflections diminish upon reducing the pH to 5.0, indicating the transition to unstructured emulsions. The graphs represent measurements from three independent experiments.

**Supporting figure 2.**
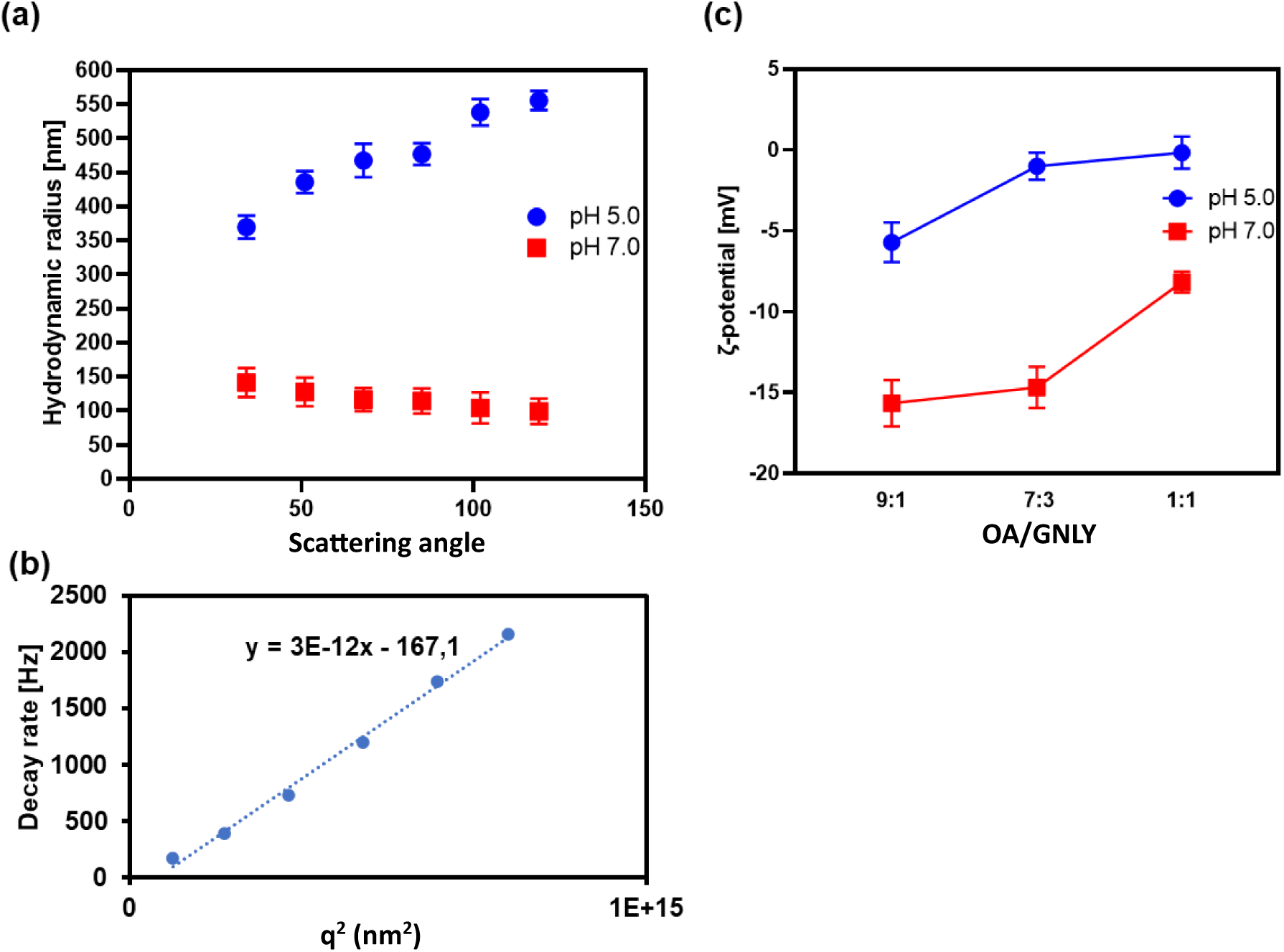
Confirmation of stable OAGNLY self-assemblies at pH 7 by DLS and electrophoretic mobility analyses. **(a)** DLS analysis reveals the hydrodynamic radius (RH) of the nanocarriers at pH 5.0 and 7.0. **(b)** Decay rate analysis of OAGNLY nanocarriers at pH 7.0 demonstrates the particles are relatively monodispersed. **(c)** ζ-potential measurements of OAGNLY nanocarriers at ratios 9:1, 7:3, and 1:1, at both pH 5.0 and 7.0, highlight the surface charge variations in different formulations.

**Supplementary figure 3.**
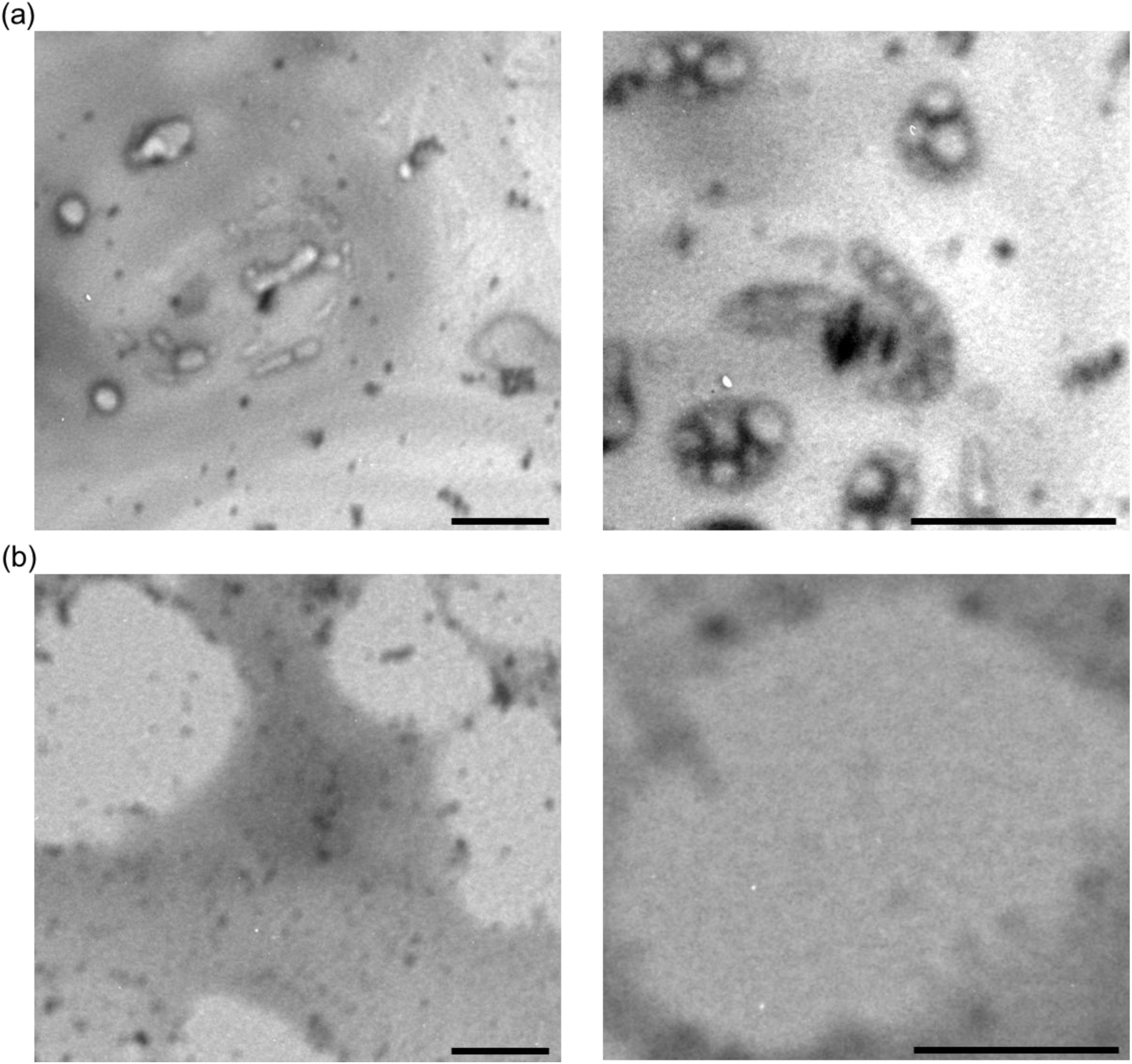
Transmission Electron Microscopy (TEM) analysis of negatively stained OAGNLY nanocarriers at different pH values. **(a)** At pH 7.0, OAGNLY forms round to rod-shaped structures of 40 to 150 nm. **(b)** At pH 5.0, OAGNLY nanocarriers display larger, unstructured arrangements, consistent with emulsions droplets of 300 to 500 nm, indicative of disrupted self-assembly under acidic conditions. Scale bars are 200 nm.

**Supporting figure 4.**
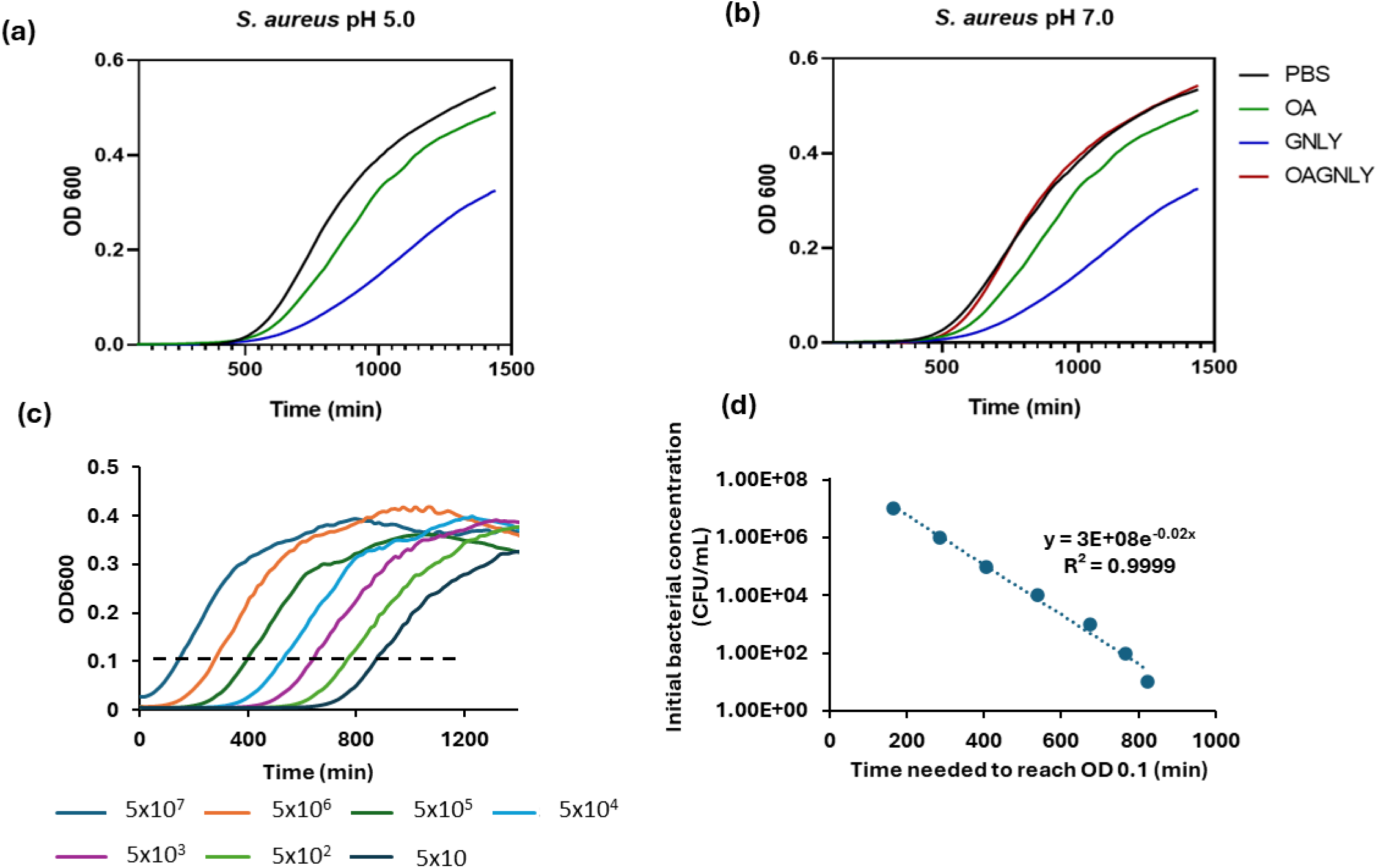
Growth curves and CFU calculations following in vitro treatment of S. aureus. 40 μL S. aureus suspensions were treated with indicated reagents in PBS for 1 hour at 37°C. After treatment, 160 μL of broth was added, and bacterial growth was monitored in a plate reader (BioTek Synergy H1) with intermittent shaking (15-second intervals every 15 minutes) for 24 hours at OD600 nm. Representative growth curves of S. aureus are shown that were treated with OAGNLY nanocarriers (32 µg/mL) or with the single components (OA 32 µg/mL, GNLY 32 µg/mL) at pH 5.0 **(a)** or at pH 7.0 **(b)**. **(c)** demonstrates reference growth curves obtained from serially diluted S. aureus cultures, showing corresponding bacterial concentrations (CFUs/mL). **(d)** Growth curves were calibrated against reference curves obtained from 10-fold serial dilutions of bacteria grown for 24 hours. A plot of the time to reach an OD600 of 0.1 (dashed line in **c**) against initial bacterial concentrations is depicted that enables CFU determination in experimental settings. A trendline with corresponding equation and correlation value is indicated.

**Supplementary figure 5.**
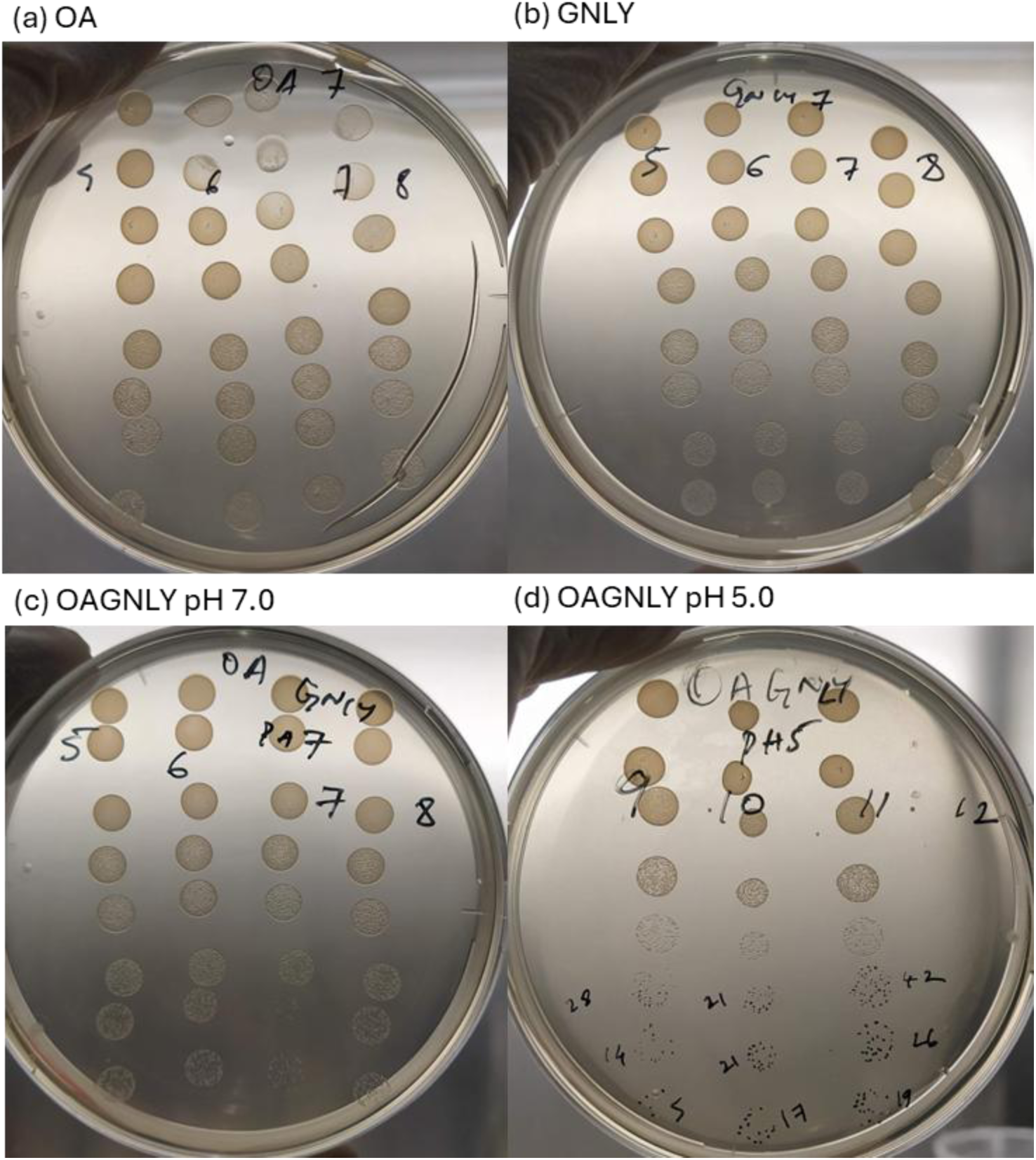
The efficacy of the treatment was assessed by spotting dilution assays. Representative results of the spotting dilution assays on agar plates after treatment of methicillin-resistant Staphylococcus aureus (MRSA) under indicated conditions. Shown are treatments with **(a)** OA 32 µg/mL alone, **(b)** GNLY 32 µg/mL alone, **(c)** OAGNLY (OA 16 µg/mL, GNLY 16 µg/mL), all at pH 7, and **(d)** OAGNLY (OA 16 µg/mL, GNLY 16 µg/mL) at pH 5.

**Supporting figure 6.**
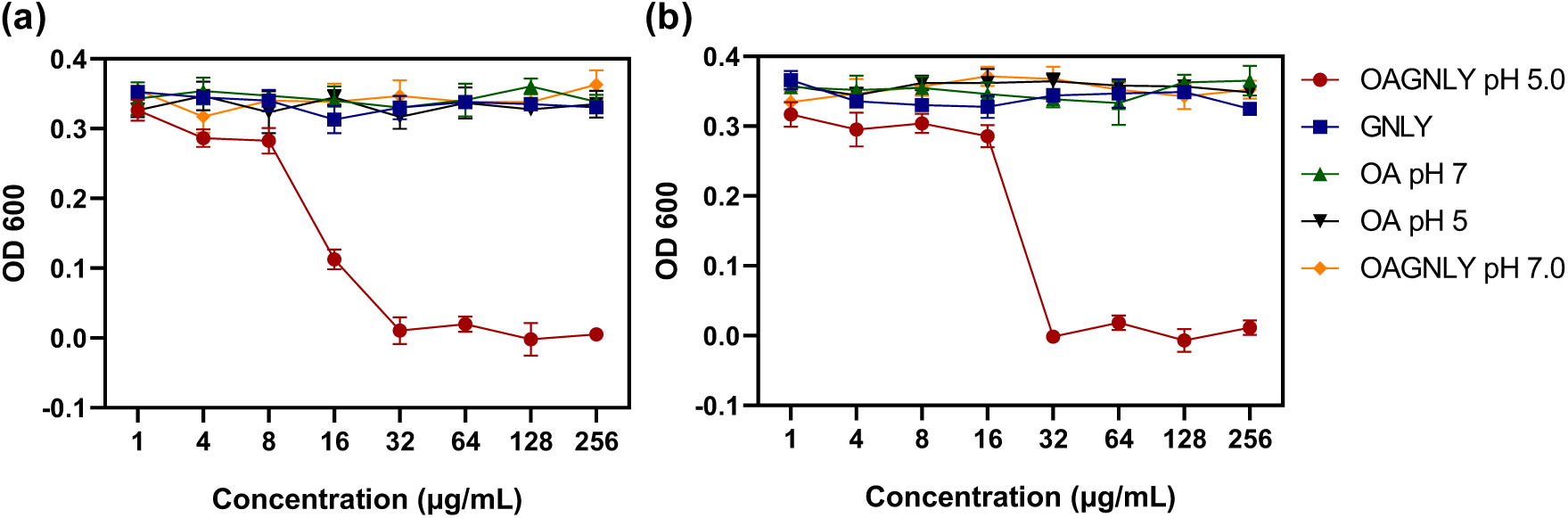
Minimal inhibitory concentration (MIC) analysis of OAGNLY nanocarriers under physiological salt conditions. **(a)** MIC values against MRSA and **(b)** CREC at pH 5.0 and 7.0 indicated by OD600 readings after 24 hours of incubation. The graphs represent data from three independent experiments, each conducted in triplicates.

**Supplementary figure 7:**
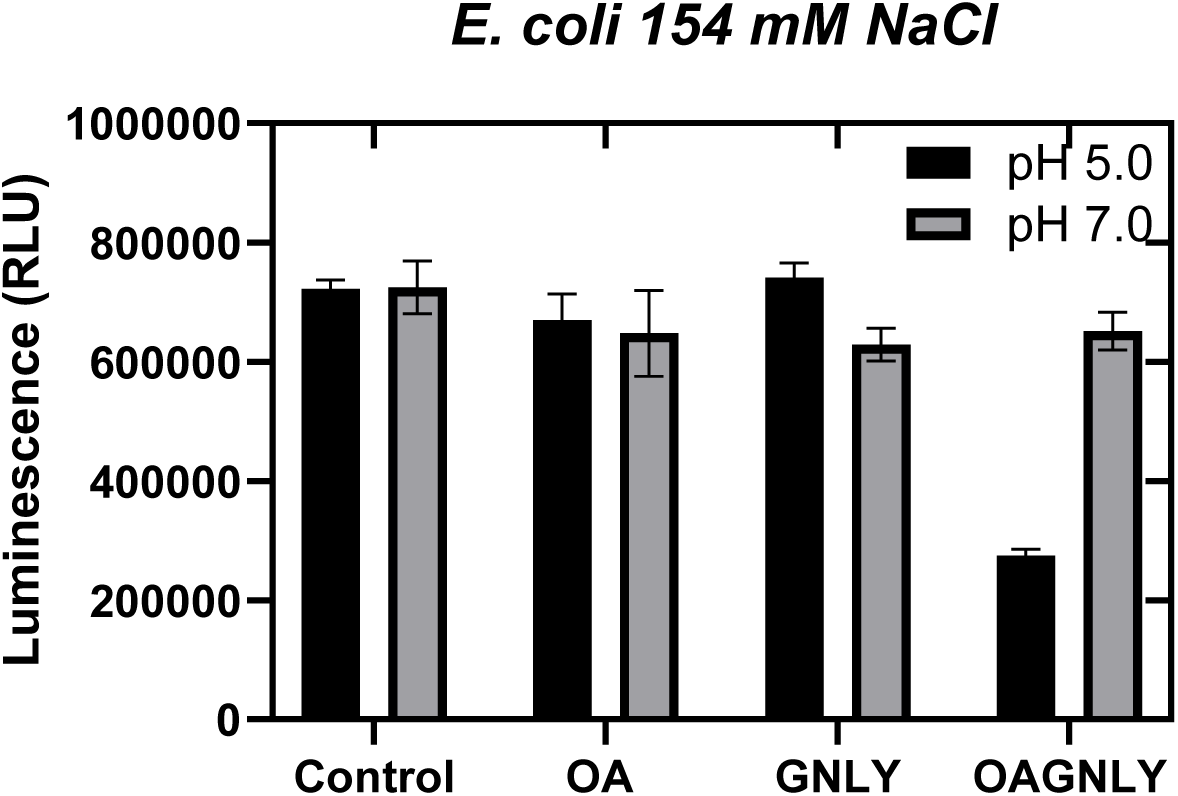
microbial cell viability assessed by ATP measurements. Bactericidal activity of OAGNLY nanocarriers (32 µg/mL) compared to their individual components (OA 32 µg/mL or GNLY 32 µg/mL) and PBS treated control samples was monitored by the BacTiter-Glo Microbial Cell Viability Assay was used to measure the luminescence, which is proportional to the amount of ATP present and indicative of metabolically active bacterial cells. The data show that treatment with OAGNLY resulted in significantly reduced luminescence, confirming the results of growth and spot dilution CFU assays. Bars represent the mean ± SEM from three independent experiments, each performed in triplicate.

**Supplementary figure 8.**
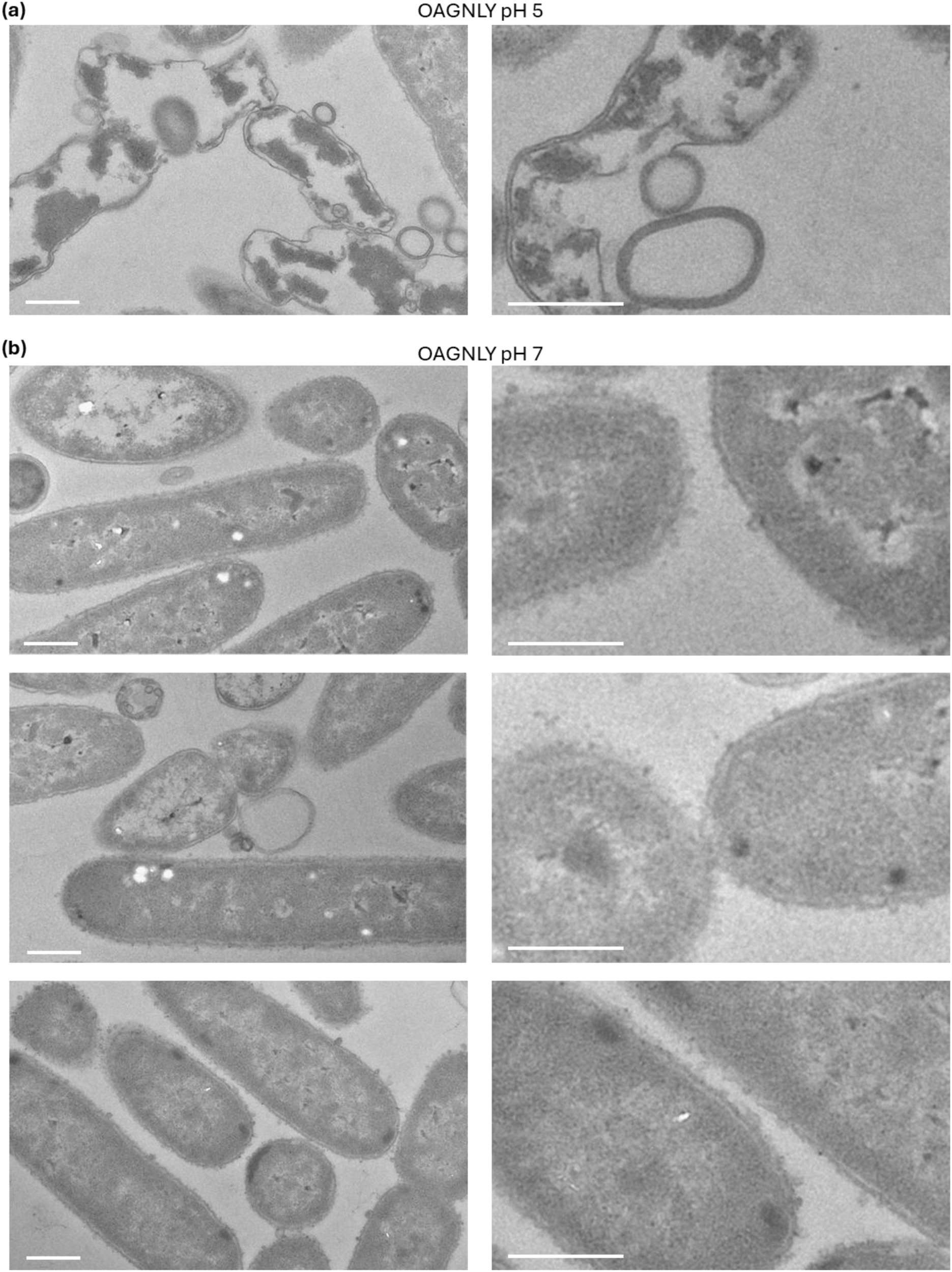
TEM analysis of E. coli morphological changes upon OAGNLY treatment. Representative images are shown for E. coli treated with OAGNLY nanocarriers for 1 hour at **(a)** pH 5 or **(b)** pH 7. After treatment at pH 7 small structures appear attached to the surfaces of otherwise unchanged bacteria. Scale bars = 100 nm. Representative images from three independent experiments are shown.

**Supplementary figure 9.**
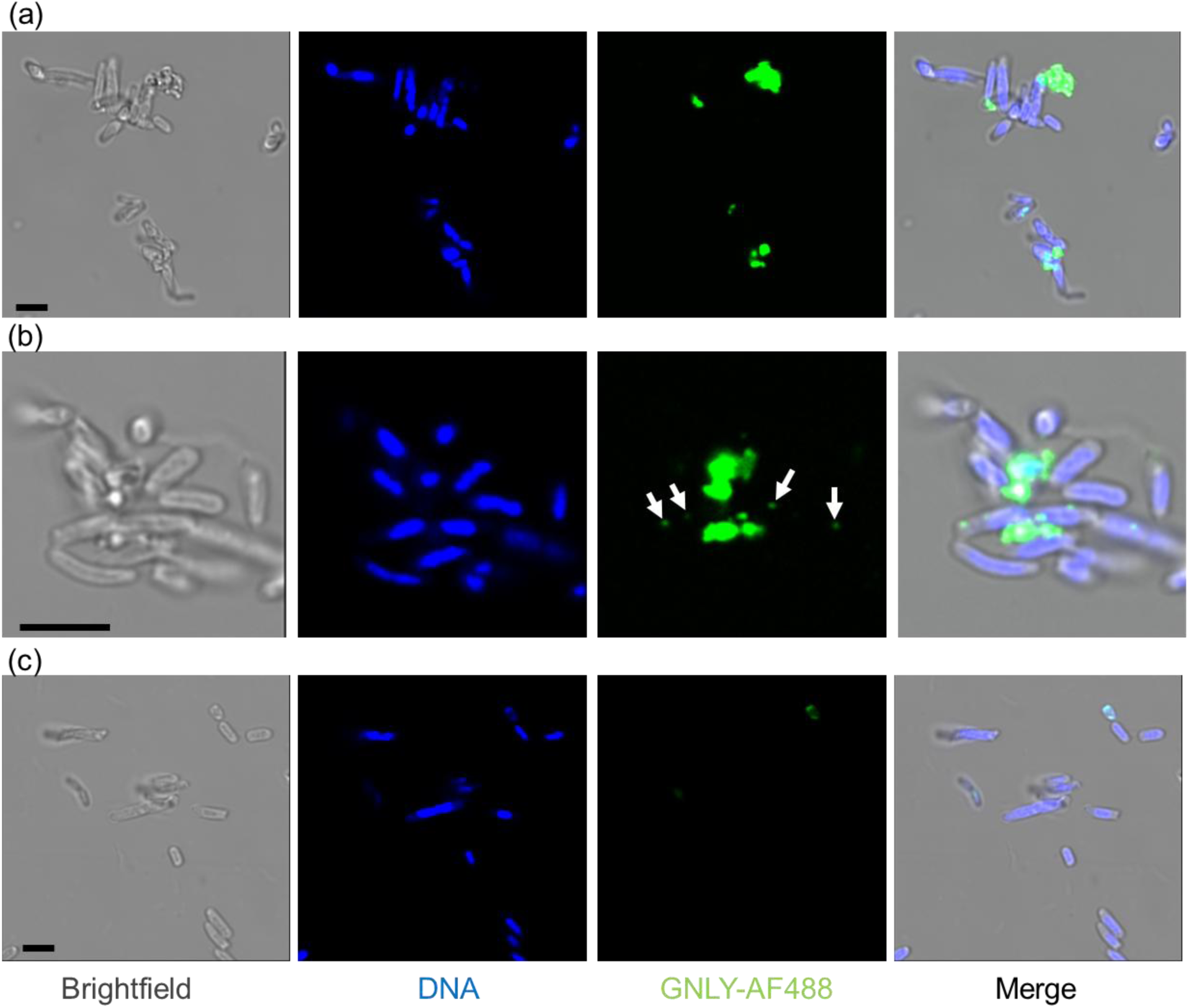
High resolution confocal microscopy imaging of OAGNLY interactions with E. coli. OA self-assemblies were generated with AF488 labeled GNLY. E. coli were treated with the fluorescent OAGNLY nanocarriers (32 µg/mL) **(a and b)** or with GNLY-AF488 alone (32 µg/mL) **(c)** for 30 min in PBS before multiple washes and PFA fixation. The specimens were counterstained with DAPI for chromosomal DNA labeling and assessed by high resolution confocal microscopy. Depicted are representative images from repeated experiments. Arrows in **(b)** show presumed single nanocarrier binding. Bars are 2 μm.

**Supplementary figure 10.**
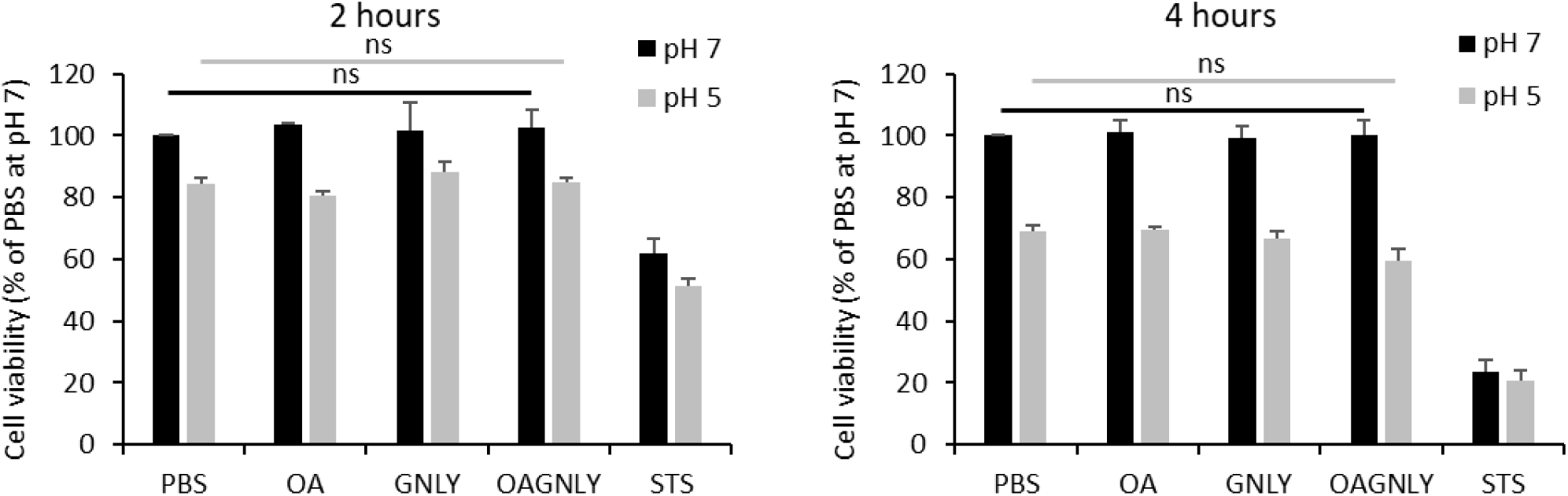
Toxicity of OAGNLY against mammalian cells at PH 5 and pH 7. HeLa cells were treated with OAGNLY 32 µg/ml (16 µg/mL OA, 16 µg/mL GNLY) or single components (32 µg/mL OA or 32 µg/mL GNLY) for 2 or 4 hours at indicated pH in PBS. Viability was assessed by MTS assay. Bars represent the mean percentage of cell viability in PBS at pH 7 ± STD from three independent experiments.

**Supporting figure 11.**
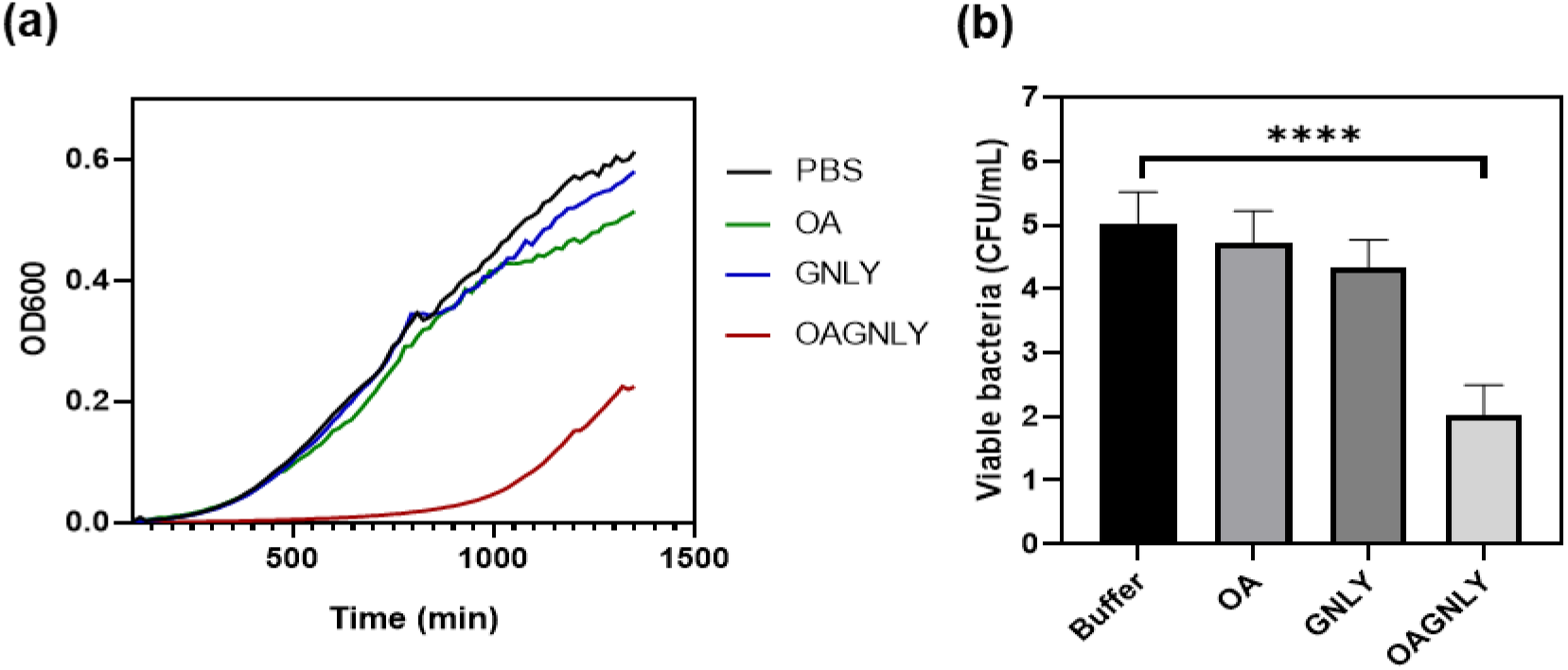
Bacterial growth and load quantification in skin homogenates after infection and treatment. **(a)** Balb/c mice were infected with MRSA contaminated sutures for 1 hour and then treated with OAGNLY (32 µg/mL) or individual components (OA = 32 µg/mL or GNLY = 32 µg/mL) for 4 hours in PBS. MRSA bacterial load in excised and homogenized surgical site wounds was quantified by growth curves analyses. Bars display mean CFUs ± SEM based on three independent experiments (n = 5 mice per group). **(b)** Quantification of bacterial loads from the same experiments, showing mean and SEM of CFUs.

**Supporting figure 12.**
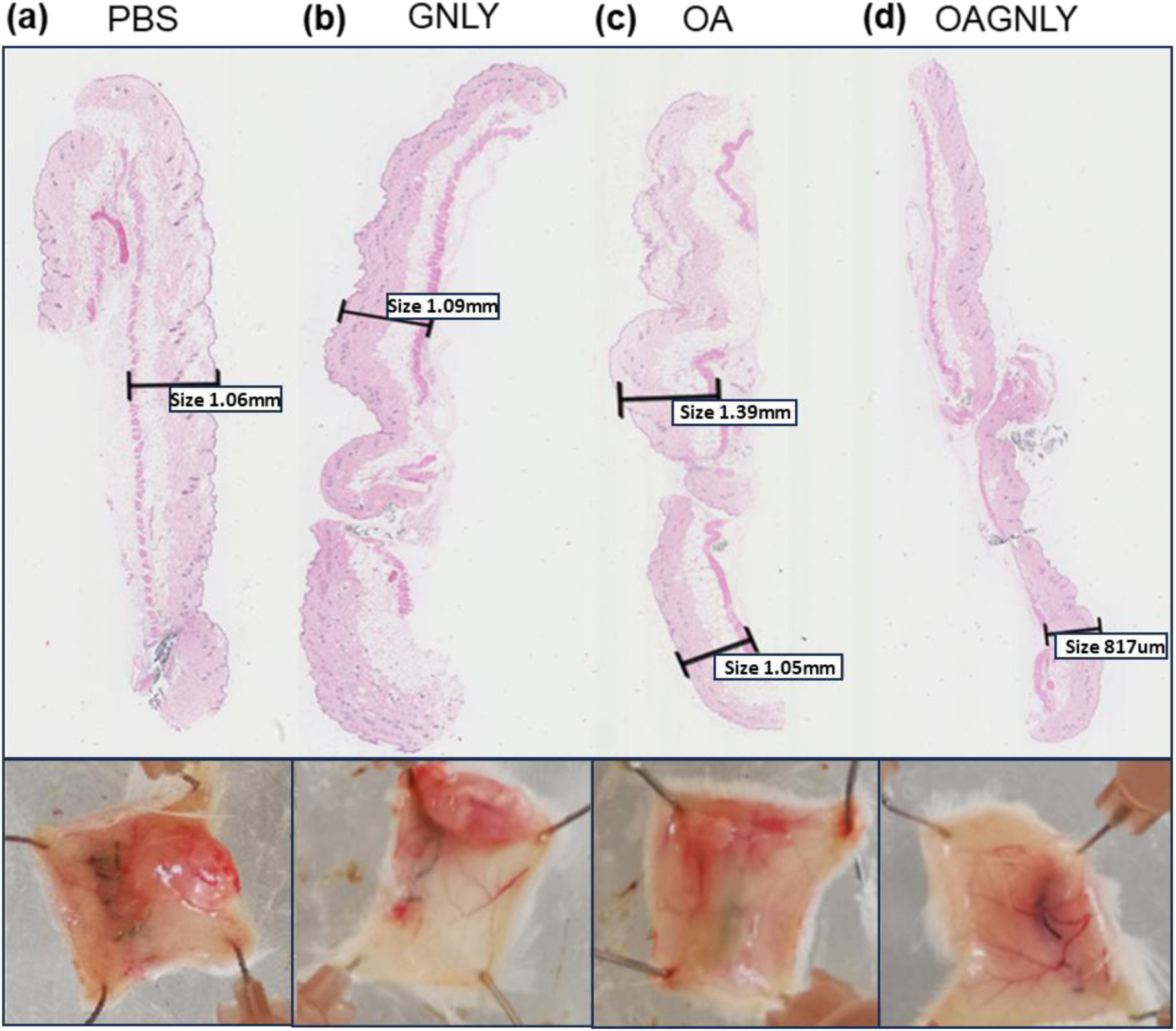
Histological analysis of skin sections from surgical site infection models. Representative H&E-stained sections show epidermal thickness (indicated) and other inflammation signs in infected skin treated with **(a)** PBS, **(b)** GNLY, **(c)** OA and **(d)** OAGNLY. Macroscopic images of skin biopsies corresponding to the same treatment conditions. Data represent three independent experiments.

**Supporting figure 13.**
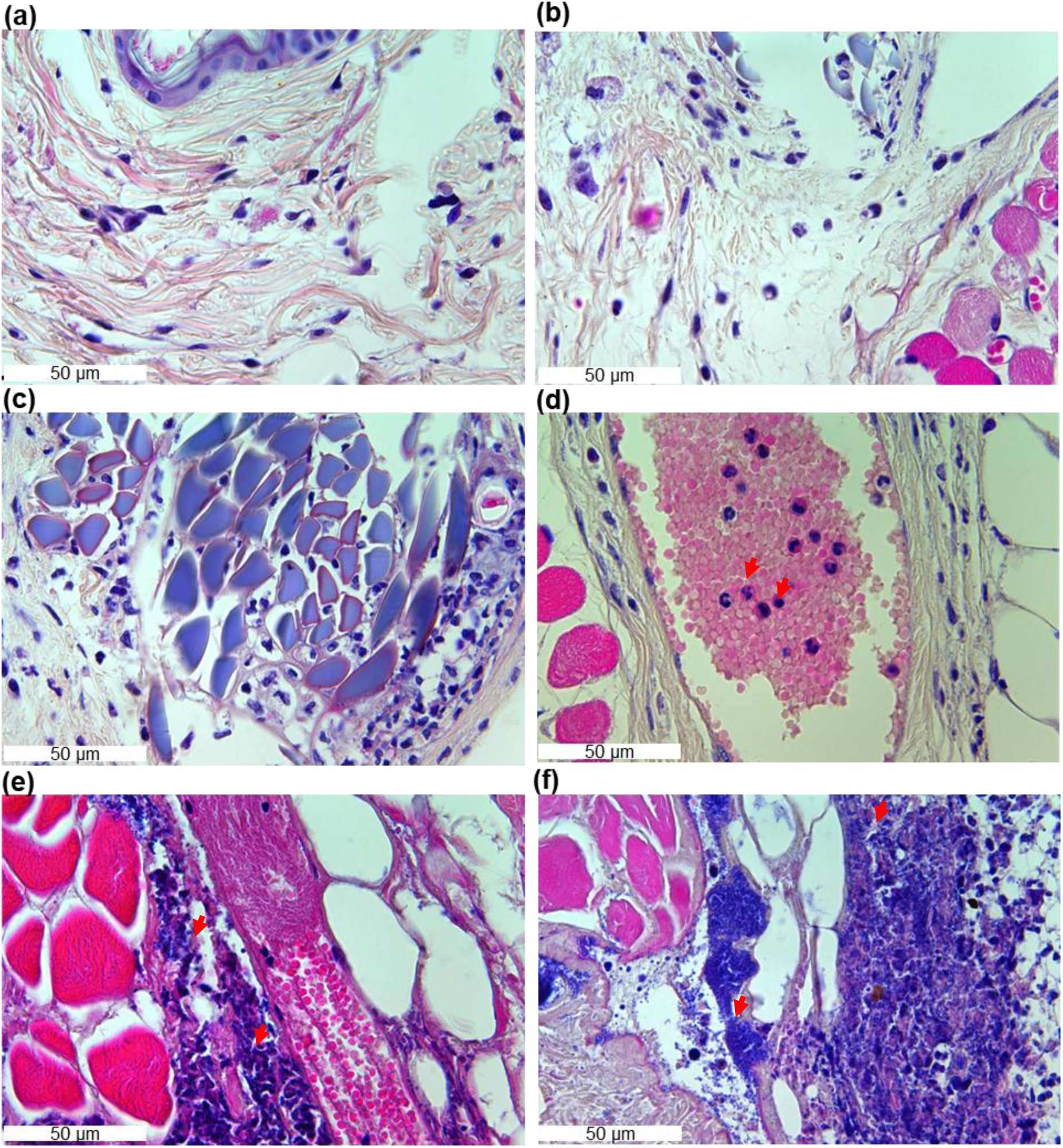
Inflammation score criteria in surgical wound models. **(a-b)** Non-infected surgical wounds show minimal inflammation compared to **(c-f)** infected surgical wounds treated with OAGNLY **(c)** or control treatments **(d-f)**. Inflammation indicators include **(d)** dilation and neutrophil sequestration in microvasculature (red arrows), **(e)** tissue infiltration of inflammatory cells (red arrows), and **(f)** bacterial invasion depth and size in surrounding tissues (red arrows). Representative images from three independent experiments are shown.

## Notes

### Competing Interest Statement

The authors have declared no competing interest.

